# Stress-induced phase separation of ERES components into Sec bodies precedes ER exit inhibition in mammalian cells

**DOI:** 10.1101/2022.02.11.480147

**Authors:** Wessel van Leeuwen, Dan T.M. Nguyen, Rianne Grond, Tineke Veenendaal, Ginny G. Farías, Catherine Rabouille

## Abstract

Phase separation of ER-exit-sites (ERES) components into membraneless compartments, the Sec bodies, occurs in *Drosophila* cells upon specific cellular stressors, i.e., salt stress and amino acid starvation, and their formation is linked to the inhibition of the early secretory pathway. Here, we show Sec bodies also form in secretory mammalian INS-1 cells upon the same stress. These reversible and membraneless structures are positive for ERES components, including both isoforms of Sec16 (A and B) and COPII subunits. We find that Sec16A, but not Sec16B, is a driver for Sec body formation. We show that the coalescence of ERES components into Sec bodies occurs by fusion, in line with their liquid-droplet properties. Lastly, we demonstrate that stress-induced ER-exit inhibition is a consequence of the significant coalescence of Sec16A into Sec bodies, leading to its depletion from ERES that become non-functional. Stress relief causes an immediate dissolution of Sec bodies and the concomitant restoration of protein exit from the ER. We propose a model in which dynamic conversion between ERES and Sec body assembly, driven by Sec16A, regulates protein exit from the ER during stress and upon stress relief in mammalian cells, thus providing a conserved pro-survival mechanism in response to stress.

## Introduction

Phase separation is an important aspect of cellular organization. It is the result of demixing or coalescence of seemingly diffuse macromolecules into non-membrane-bound compartments. Phase separation occurs both in the cytoplasm and in the nucleus where it separates two distinct phases (Gomes and Shorter, 2018; Hyman et al., 2014). Interestingly, phase separation is often initiated and driven by ‘’driver’’ proteins that engage each other by low affinity multivalent interactions (Banani et al., 2017). These driver proteins contain intrinsically disordered low complexity domains that comprises repeating sequences and low amino acid variation (Franzmann and Alberti, 2019; Martin and Mittag, 2018). Absence of driver proteins leads to unstable phase separated compartments or they will not form.

Phase separation can also be driven by stress (such as oxidative, ER stress, heat shock and nutrient starvation), resulting in the formation of stress assemblies (van Leeuwen and Rabouille, 2019). One of the most studied stress assemblies are P-bodies and stress granules that form around RNAs (Jain and Parker, 2013; Wheeler et al., 2016). Recently, a new stress-driven phase separated compartment has been identified in *Drosophila* S2 cells, namely the Sec body (Zacharogianni et al., 2014).

Sec bodies are related to the early secretory pathway, a major anabolic pathway used by 30% of proteins to reach their functional localizations (Sharpe et al., 2010). They form at the Endoplasmic Reticulum exit sites (ERES), ribosome-free regions of the ER where newly synthesized proteins destined to nearly all membrane compartments of the cell as well as to the extracellular space, exit via COPII coated vesicles. In S2 cells, Sec bodies contain COPII subunits that form the COPII coat itself, the inner coat proteins Sec23/24 and the outer coat proteins Sec13/31 (Miller and Schekman, 2013). They also contain the ERES large peripheral scaffold protein Sec16, a key regulator in the organization of the ERES and in COPII coated vesicle budding (Sprangers and Rabouille, 2015).

In *Drosophila* S2 cells, Sec bodies form upon specific stress that drive these ERES components (Sec16, Sec23 and Sec31) to coalescence into micrometer large membraneless assemblies. Sec bodies form as 1-5 rounded structures per cell (Zacharogianni et al., 2014; Zhang et al., 2021), and their diameter ranges from 0.4 to 1 μm. They are membrane-less, polyadenylated RNA-free, and display liquid-like properties (Zacharogianni et al., 2014). Sec bodies are reversible upon stress removal, and they contribute to cell survival (Aguilera-Gomez et al., 2016; Zacharogianni et al., 2014). In addition, Sec16 has been shown to be a driver for Sec bodies formation in S2 cells. Its depletion prevents Sec body formation and overexpression of a conserved 44 amino-acid C-terminal domain of Sec16, SRDC (<u>S</u>erum <u>R</u>esponsive <u>D</u>omain <u>C</u>onserved), drives Sec body formation, even in the absence of stress (Aguilera-Gomez et al., 2016).

Although Sec bodies are well characterized in *Drosophila* cells, their formation has remained elusive in mammalian cells. In this study, we took advantage of the knowledge gained from *Drosophila* S2 cells to investigate Sec body formation in mammalian cells. Here, we found that highly secretory cells, such as INS-1 cells (rat pancreatic insulin secreting cells that are widely used as rat islet β cell model), form Sec bodies similar to *Drosophila* S2 cells. Indeed, using immunofluorescence for endogenous Sec16A, we found that this protein coalesces into large and rounded structures upon high salt stress and amino acid starvation in KRBm, that is a Krebs Ringer Bicarbonate buffer that we adapted for mammalian cells. These structures also contain COPII subunits, are reversible upon stress relief, and have features of liquid droplets. Using immuno-electron microscopy (IEM), we also show that they are membrane-less but in very close proximity to the ER. All these features clearly identify these structures as Sec bodies. Furthermore, we found that, from the two Sec16 encoded proteins in mammalian cells, Sec16A is a clear driver in their formation. Indeed, knockdown of Sec16A, but not Sec16B, prevents Sec body formation upon stress, and its sole overexpression accelerates stress-induced Sec body formation. Using live-cell imaging, we show that Sec16A and Sec16B are simultaneously recruited into the same Sec body during early assembly, along COPII subunits. Live-cell imaging also revealed that Sec bodies form by fusion of remodeled ERES. Finally, applying the RUSH (retention using a selective hook) transport assay, we found that the inhibition of ER exit is a consequence of the formation of Sec bodies that recruits Sec16A/B and COPII proteins, progressively depleting the cells of functional ERES. Immediate dissolution of Sec bodies followed by restored protein ER exit was observed upon stress removal. Our findings suggest that dynamic assembly and disassembly of Sec bodies regulates protein secretion in mammalian cells during periods of stress and stress relief.

## Results

### Upon NaCl stress and KRBm incubation, the ERES of INS-1 cells are remodeled into large Sec16A positive structures resembling Sec bodies

Recent work in *Drosophila* S2 cells has shown that Sec body formation is promoted by two specific types of stress. It is triggered by high salt stress (addition of NaCl to the growing medium) and by incubating the cells in Krebs Ringer Bicarbonate buffer (KRB) leading to a moderate salt stress that is potentiated by the absence of amino acids (Zacharogianni et al., 2014; Zhang et al., 2021).

To assess whether Sec bodies also form in mammalian cells, we first incubated a range of mammalian cell lines, including HepG2, MDCKII and MRC5, upon addition of 250 mM NaCl for 4 h to their growing medium. This produced some remodeling of their ERES. However, none was as pronounced as in S2 cells and occurred only in a low percentage of cells (Figure 1—figure supplement 1A). We then assessed rat insulinoma INS-1 cells that secrete insulin and appear to have an active metabolism (Cline et al., 2011). Incubation of INS-1 cells in RPMI medium supplemented with 200 mM NaCl (RPMI200) led to a remarkable remodeling of ERES that was visualized by immunofluorescence using a specific antibody against endogenous Sec16A. In basal conditions, Sec16A localizes at ERES (Hughes et al., 2009; Tillmann et al., 2015; Watson et al., 2006), but upon high NaCl, it coalescences into round and bright remodeled structures that resemble S2 cell Sec bodies (**Figure 1A**).

**Figure 1.**
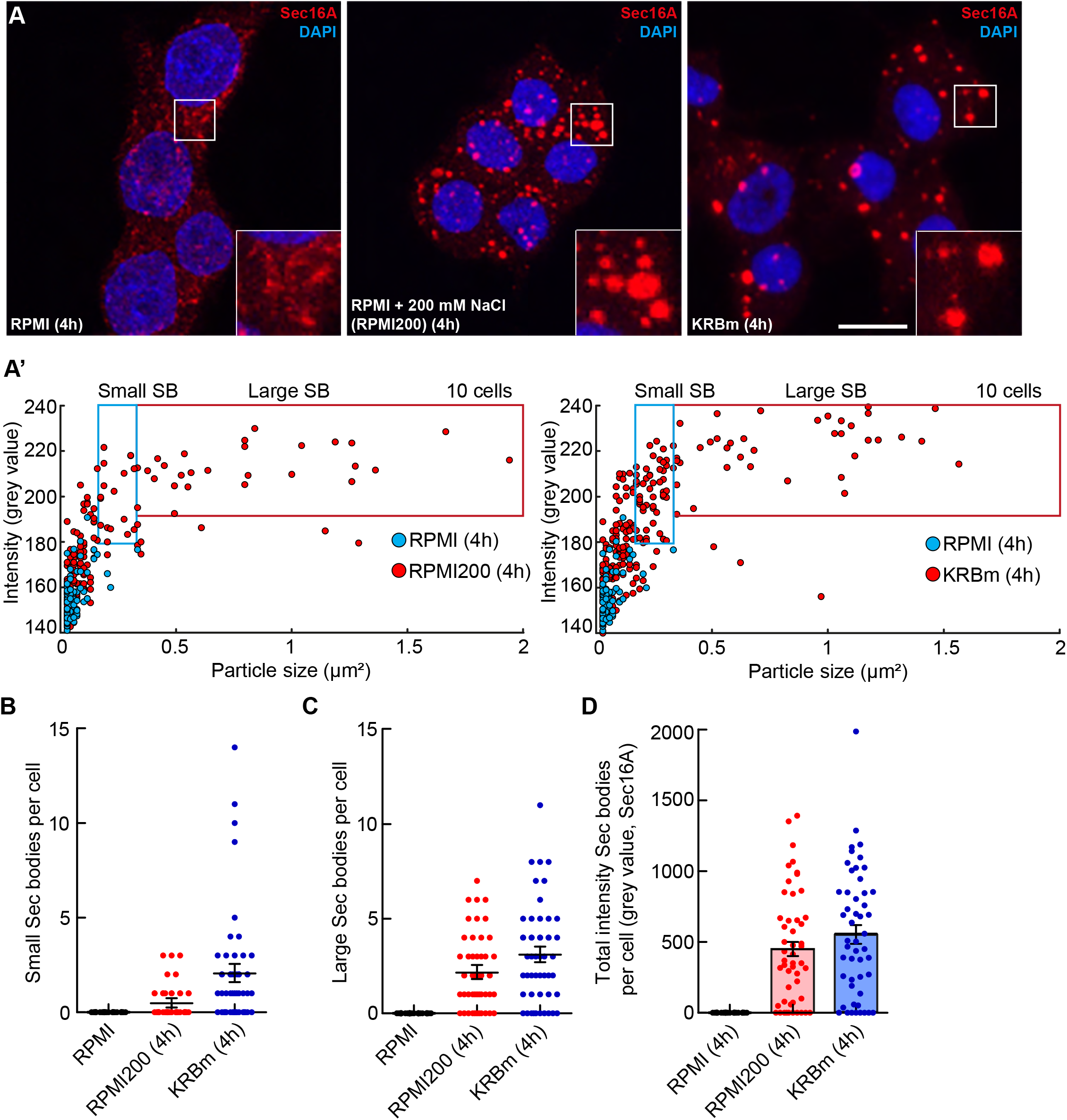
Remodeling of ERES into large structures resembling Sec bodies in INS-1 cells upon salt stress and KRBm. **A:** Immunofluorescence visualization of endogenous Sec16A (red) in INS-1 cells upon incubation in RPMI, RPMI200 (RPMI+200 mM NaCl) and KRBm (4 h). Note that, upon stress, Sec16A remodels into small and large round structures that resemble S2 cells Sec bodies. **A’:** Scatterplot depicting Sec16A foci size and intensity upon incubation in RPMI, RPMI200 and KRBm (4 h). All foci in the red box are determined as ‘’large Sec bodies’’ and foci in the blue box are ‘’small Sec bodies’’. The values within the red and blue box were used to determine the total intensity of Sec bodies per cell and per condition. Foci from a total of 10 cells are displayed. **B:** Dot plot depicting the number of small Sec16A-positive Sec bodies per cell in INS-1 cells upon RPMI, RPMI200 and KRBm (4 h); n=50 cells. **C:** Dot plot depicting the number of large Sec16A-positive Sec bodies per cell in INS-1 cells upon RPMI, RPMI200 and KRBm (4 h); n=50 cells. **D:** Dot and bar plot depicting the total intensity of Sec16A-positive Sec bodies per cell in INS-1 cells upon RPMI, RPMI200 and KRBm (4 h). n=50 cells. See also Figure 1—figure supplement 1. Scale bar: 10 μm (A). Error bar: SEM (B-D).

We then designed a method to assess the efficiency of Sec16A remodeling in INS-1 cells considering the number, size (area) and mean intensity of coalesced structures (**Figure 1A’**; see also Material & Methods). Accordingly, we found that RPMI200 leads to the formation of an average of 0.5 small (0.15-0.3 μm^2^), and 2.2 large (>0.3 μm^2^) remodeled round and bright structures per cell (**Figure 1A’-C**). By multiplying the size and mean intensity of each small and large structure, the remodeling can also be represented as the total intensity of all remodeled Sec16A positive structures per cell. This gives a clear read-out of remodeling efficiency (i.e., Sec body formation, see below) upon stress (**Figure 1D**). Upon addition of NaCl, Sec16A remodels into large structures in a time dependent manner, in which 4 h triggers a more efficient remodeling, when compared to shorter incubation periods (Figure 1—figure supplement 1B). These large structures are very reminiscent of Sec bodies observed in *Drosophila* S2 cells, and they are present in 72 % of the incubated cells (**Figure 1D**).

As in S2 cells, the formation of these structures in INS-1 cells is specific for addition of NaCl, as addition of 200 mM either NaAcetate or KCl to RPMI does not lead to the formation of large circular Sec16A positive structures. Neither does the addition of 0.4 M sorbitol, indicating that osmotic shock is not involved in this response (Figure 1—figure supplement 1C, C’).

In parallel, a 4 h incubation of INS-1 cells in KRBm (KRB buffer for mammalian cells) which induces a moderate salt stress combined with amino-acid starvation) also leads to the efficient formation of Sec16A positive remodeled structures (**Figure 1A, A’**). Cells displayed an average of 2.1 small and 3.1 large, remodeled structures per cell, leading to an increase in total intensity of remodeled Sec body-like structures per cell similarly to high NaCl (**Figure 1B-D**). Large Sec16A structures are present in 78% of the incubated cells (**Figure 1D**).

Overall, this data indicates that as in S2 cells, both NaCl stress and amino acid starvation in KRBm leads to the remodeling of ERES of INS-1 cells into structures that morphologically resemble Sec bodies.

### Stress-induced Sec16A positive structures are Sec bodies

To determine whether these Sec16A remodeled structures observed in INS-1 cells are indeed Sec bodies, we first assessed whether they also contain COPII subunits, a documented feature of *Drosophila* Sec bodies. Accordingly, we found that endogenous Sec13 and Sec24A co-localize with Sec16A under both RPMI200 and KRBm (**Figure 2A, B**), with a Mander’s overlap coefficient of around 0.75 for Sec16A/Sec13 (**Figure 2A’**) and 0.76 for Sec16A/Sec24A (**Figure 2B’**) under both stress conditions. In addition, we investigated whether these stress-induced structures contain other proteins of the early secretory pathway. p115, a protein that facilitates COPII vesicle transport from the ER to the Golgi apparatus (Sapperstein et al., 1995; Waters et al., 1992), partially co-localized with Sec16A in remodeled structures (Figure 2—figure supplement 1A), but not GM130, GRASP65 and GRASP55 (Figure 2—figure supplement 1B-D). Overall, the remodeling of Sec16A in INS-1 cells does not appear to incorporate a large set of Golgi proteins.

**Figure 2.**
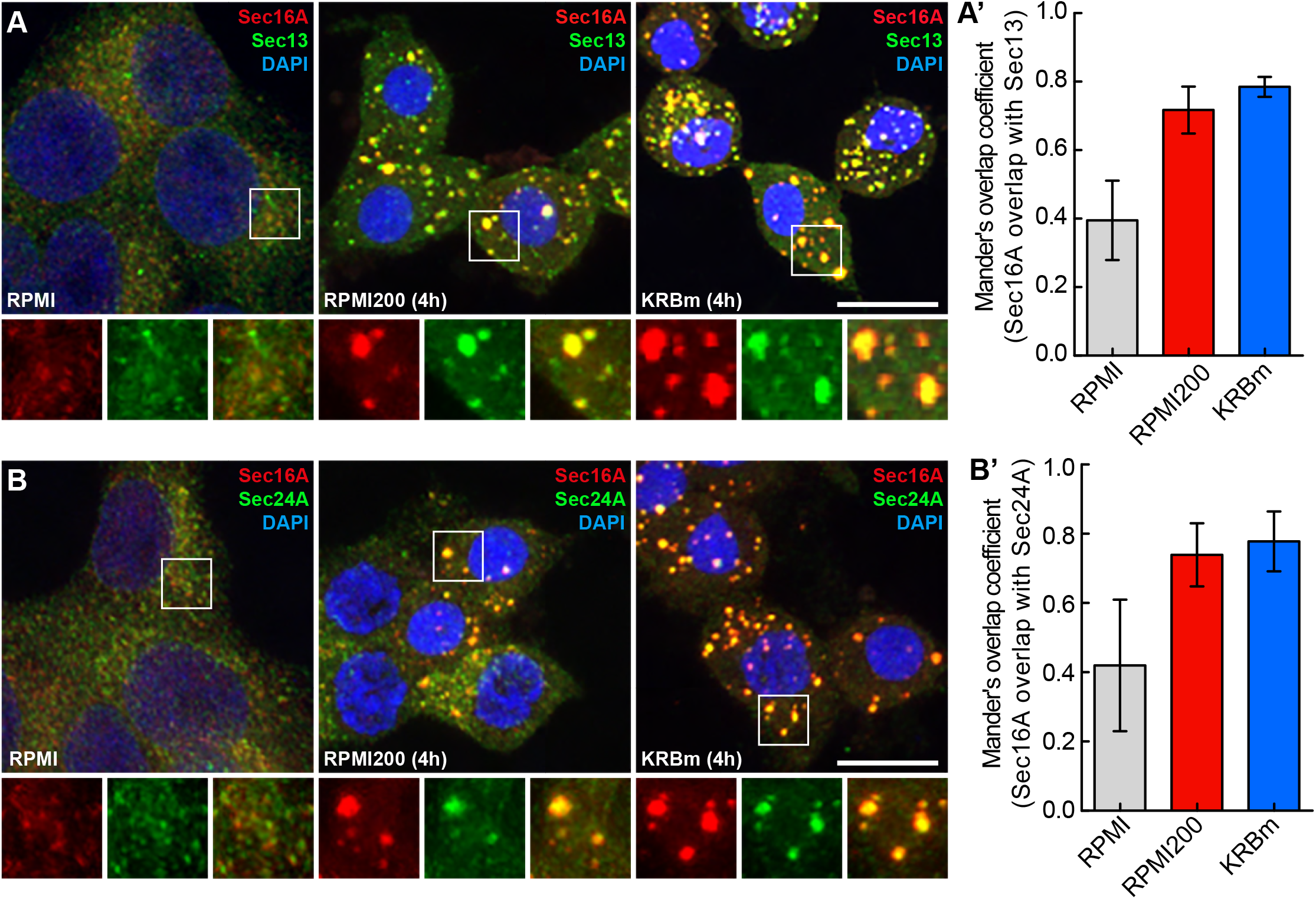
Sec16A-positive structures contain COPII subunits. **A, A’:** Immunofluorescence visualization of endogenous Sec16A and Sec13A in INS-1 cells upon incubation in RPMI, RPMI200 and KRBm (4 h) (A). Quantification of the Mander’s overlap coefficient between Sec16A and Sec13 (A’); n=2 experiments, both exp > 30 cells **B, B’:** Immunofluorescence visualization of Sec16A and Sec24A in INS-1 cells upon incubation in RPMI, RPMI200 and KRBm (4 h) (B). Quantification of the Mander’s overlap coefficient between Sec16A and Sec24A (B’); n=2 experiments, both exp > 30 cells. See also Figure 2—figure supplement 1. Scale bar: 10 μm (A) and (B) Error bar: SEM (A’) and (B’)

A second critical feature of Sec bodies in S2 cells is that they are not enclosed by a sealed lipid membrane (Zacharogianni et al., 2014). To unravel this feature in stressed INS-1 cells, we employed immuno-electron microscopy (IEM) to visualize their morphology after labelling with an antibody against endogenous Sec13. Either upon RPMI200 or KRBm, Sec13 positive coalescences appear as lightly electron dense structures with a diameter between 0.3-1 μm. Consistent with being a “membrane-less” assembly, the Sec13 positive structures are not surrounded by membrane, but often position in close proximity to the ER membrane (**Figure 3A**).

**Figure 3.**
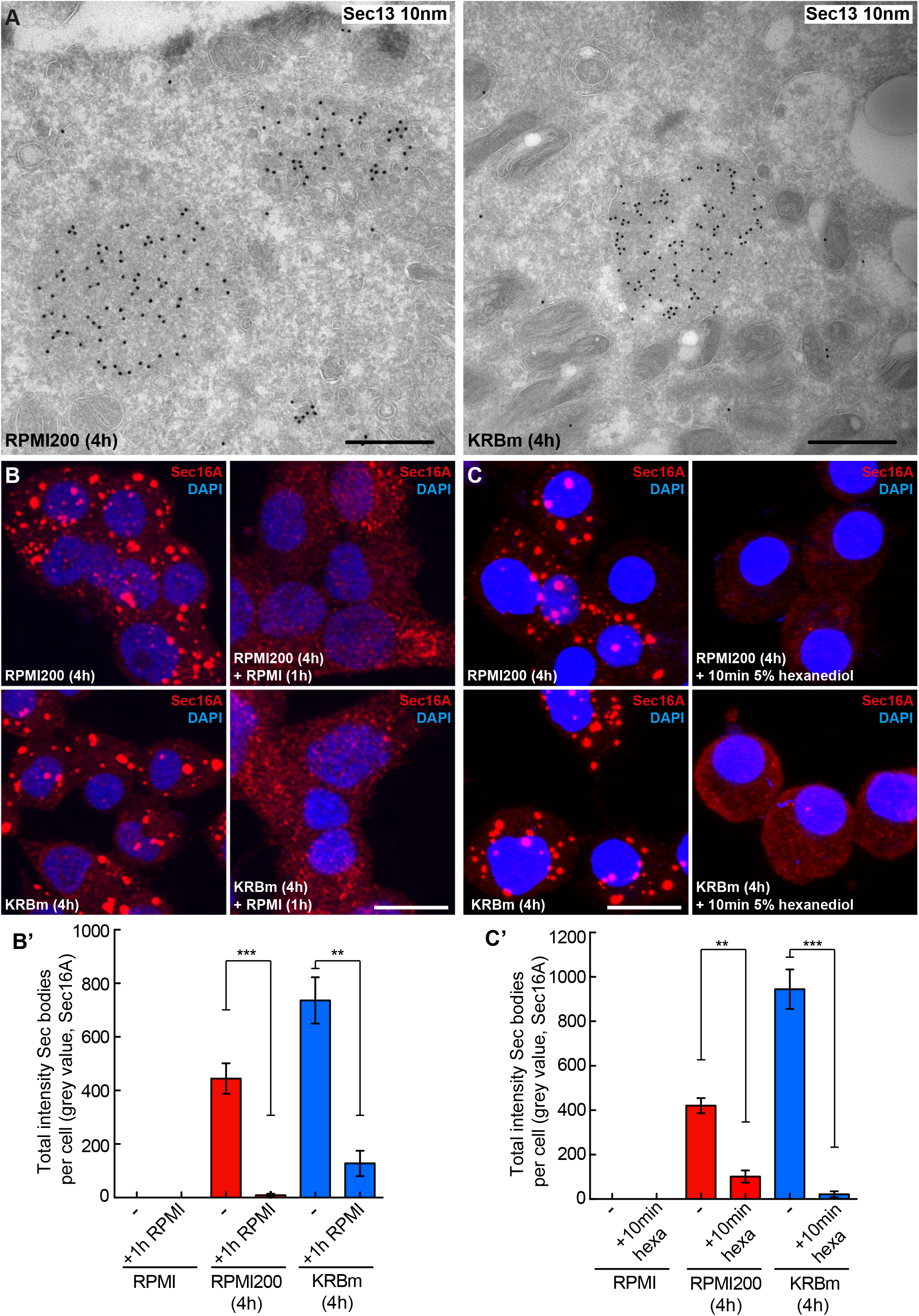
Sec16A-positive structures in stressed INS-1 cells are Sec bodies. **A, A’:** Visualization of endogenous Sec13 (10 nm cPAG) by immuno-electron microscopy in ultrathin frozen sections of INS-1 cells incubated in RPMI200 and KRBm for 4 h. Sec13 is concentrated in structures (Sec bodies) that are slightly denser than the surrounding cytoplasm, round, not sealed in a lipid bilayer and in close proximity to the ER. **B, B’:** Immunofluorescence visualization of Sec16A in INS-1 cells after 4 h in RPMI200 and KRBm followed by 1 h in RPMI (B). Note that Sec body dissolution is complete within 1 h of stress removal and that Sec16A is localized again at ERES. Quantification in (B’); n=2 experiments, both exp > 30 cells. **C, C’:** Immunofluorescence visualization of Sec16A in INS-1 cells treated with 5% hexanediol for 10 min after incubation in RPMI200 or KRBm for 4 h (C). Quantification in (C’); n=2 experiments, both exp > 30 cells. Scale bar: 500 nm (A), 10 μm (B) and (C). Error bar: SEM; **p<0.01 and ***p<0.001.

Furthermore, consistent with formation by phase separation, Sec bodies in S2 cells are reversible upon stress removal. Therefore, they are not irreversible aggregates (Zacharogianni et al., 2014). In this regard, we found that the remodeled structures are largely dissolved within 1 h of stress relief (i.e., further incubation in RPMI), and that Sec16A has regained its typical ERES pattern (**Figure 3B, B’)**.

Last, phase separation can lead to liquid as well as solid assemblies, both reversible (Saad et al., 2017; Wheeler et al., 2016). In S2 cells, Sec bodies were shown using FRAP to have liquid-like properties (Zacharogianni et al., 2014). To assess this feature for the Sec16A positive structures triggered by stress in INS-1 cells, we used 1,6-hexanediol, an aliphatic alcohol that has been used to differentiate between the two states of membrane less assemblies (Kroschwald et al., 2015; Peskett et al., 2018). Liquid assemblies are sensitive to this hexanediol and dissolve, whereas solid assemblies do not. Addition of 5% hexanediol for 10 min to RPMI200 and KRBm stressed INS-1 cells leads to the complete dissolution of the Sec16A positive structures (**Figure 3C, C’**), suggesting that they are liquid-like assemblies. Of note, 1,6-hexanediol also dissolves the typical ERES in non-stressed cells, suggesting that ERES themselves are also liquid like assemblies (Gallo et al., 2020). Overall, this data indicates that INS-1 remodeled Sec16A positive structures have liquid-like features.

Taken together, these results indicate that Sec16A remodeled into small and large round structures, induced by high salt stress and amino acid starvation in KRBm, correspond to *bona fide* Sec bodies. They contain COPII subunits, and they are membrane-less reversible stress assemblies with features of liquid droplets.

### Sec16A is a driver in Sec body formation

As mentioned in the introduction phase separation is often driven by proteins that contain disordered regions. The sequence of the single *Drosophila* Sec16 homologue comprises mostly disordered regions (except for the central conserved domain). We have shown that Sec16 is a driver for Sec body formation in *Drosophila* S2 cells. Its depletion prevents Sec body formation and the overexpression of a 44 amino-acid peptide (SRDC) located in the C-terminus of the protein is enough to drive Sec body formation (Aguilera-Gomez et al., 2016).

Rat Sec16A also harbors a large amount of disordered region in its sequence and contains a conserved 44 residues SRDC (Figure 4—figure supplement 1A, B) that is instrumental in the formation of Sec bodies in *Drosophila* cells (Aguilera-Gomez et al, 2016). In this respect, we tested whether it is also a driver in Sec body formation in INS-1 cells. Knockdown of Sec16A using siRNA (that specifically depletes *Sec16A* mRNA by 86 %, **Figure 4A**) leads to a strong inhibition of Sec body formation (visualized with endogenous Sec13), both upon KRBm incubation (**Figure 4B, B’)** and RPMI200 (Figure 4—figure supplement 1C, C’). The inhibition of Sec body formation is stronger in the cells where Sec16A is hardly detectable by IF (arrow in **Figure 4B**). In cells that still exhibit a small pool of Sec16A, the resulting Sec13 positive structures are much smaller than typical Sec bodies (arrowhead in **Figure 4B**). These data demonstrate that Sec16A is essential for the formation of Sec bodies in INS-1 cells.

**Figure 4.**
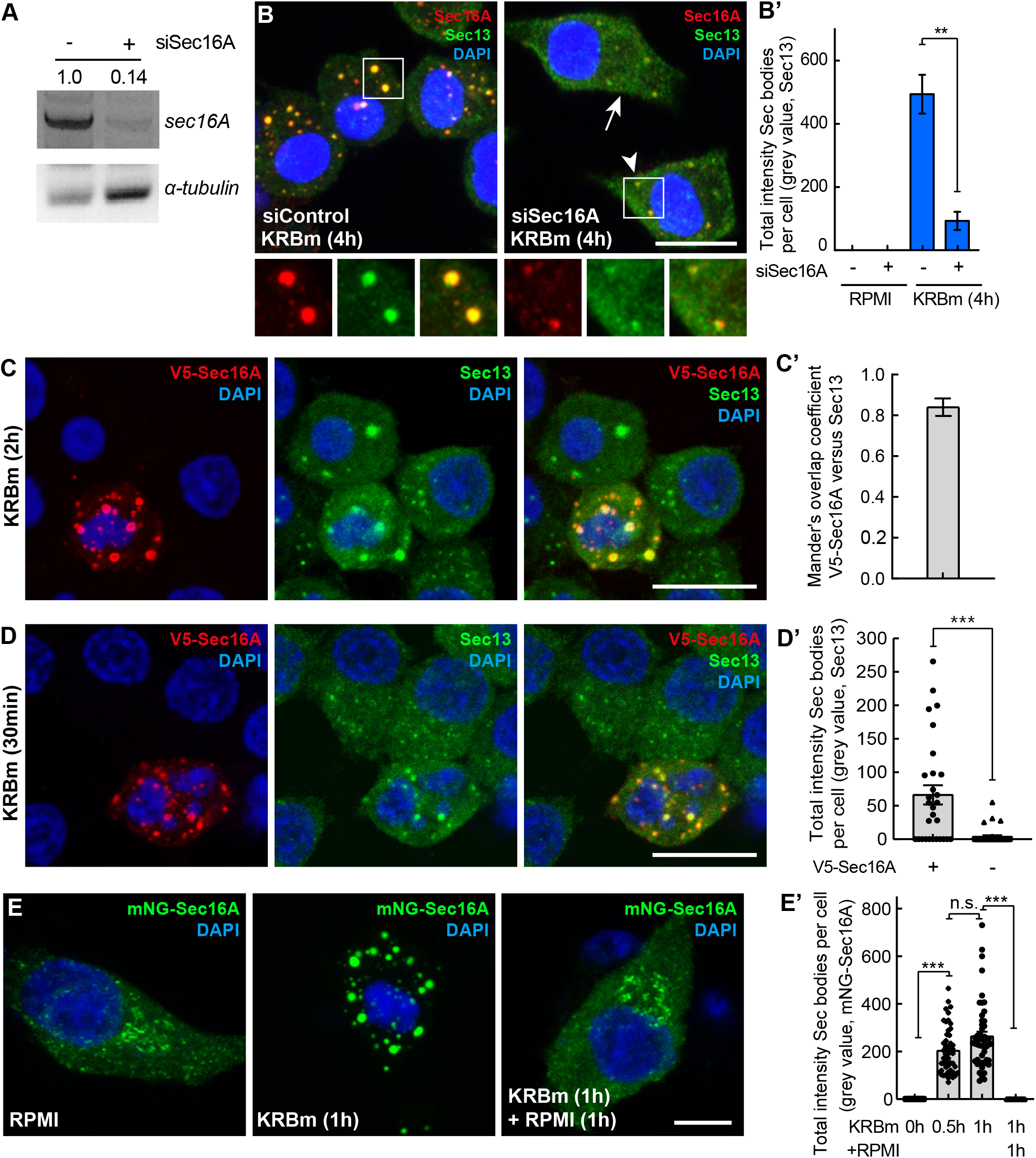
Sec16A drives Sec body formation in INS-1 cells. **A:** PCR of Sec16A and α-tubulin cDNAs from mock and Sec16A-depleted INS-1 cells. Note that depletion of Sec16A for 4 days leads to a very efficient knockdown (86%). Normalized value of Sec16A against α-tubulin, was used to calculate the Sec16A ratio between mock and Sec16A-depleted cells. **B, B’:** Immunofluorescence visualization of Sec16A and Sec13 in Sec16A depleted INS-1 cells incubated in KRBm for 4 h (B). Note that Sec body formation (marked by Sec13) is inhibited upon Sec16A depletion. Quantification of the total intensity of Sec bodies per cell (B’); n=2 experiments, both exp > 30 cells. **C, C’:** Immunofluorescence visualization of V5-Sec16A (anti V5 and endogenous Sec13 in V5-Sec16A transfected INS-1 cell after KRBm incubation for 2 h (C). Mander’s overlap coefficient between V5 and Sec13 in Sec bodies (C’); n=40 cells. **D, D’:** Immunofluorescence visualization of V5-Sec16A (anti V5) and endogenous Sec13 in V5-Sec16A transfected INS-1 cell after KRBm incubation for 30 min (D). Compare the Sec body formation in transfected versus adjacent non-transfected cells. Quantification of Sec13-positive Sec body total intensity per cell in transfected and non-transfected cells (D’); n=29 and 36 cells, respectively. **E, E’**: Fluorescence images of INS-1 cells expressing mNeonGreen (mNG)-Sec16A incubated in KRBm for 0, 60 minutes, and 60 min followed by 1h further incubation in RPMI (E). Quantification of mNG-Sec16A total intensity of Sec bodies (E’); n= 50-51 cells. See also Figure 4—figure supplement 1. Scale bars: 10 μm (B-D) and 5 μm (E). Error bar: SD (C’), SEM (B’), (D’) and (E’); ns – not significant; **p<0.01; ***p<0.001.

As Sec16A appears to be a driver in Sec body formation, we then asked whether its overexpression is also able to induce Sec body formation. Upon overexpression of V5-tagged Sec16A, we observed its localization in Sec bodies upon KRBm incubation for 2 h (**Figure 4C**). The formation of these structures is stress specific, as tagged-V5-Sec16A is localized to ERES in cells growing in RPMI (Figure 4—figure supplement 1D). Importantly, V5-Sec16A co-localizes with endogenous Sec13 (**Figure 4C**), with a Mander’s coefficient of around 0.8 (**Figure 4C’**) showing that tagged-Sec16A is incorporated into Sec bodies upon KRBm incubation for 2h.

In non-transfected cells INS-1 cells incubated with KRBm for short length of time (30 min), Sec body formation is only observed in 11% of the cells (**Figure 4D, D’,** non-transfected). We used this to ask whether the overexpression of V5-Sec16A promotes Sec body formation in this short period of stress. Interestingly, we found that V5-Sec16A overexpression leads to Sec13-positive Sec body formation in 62% of the cells, already after 30-min incubation in KRBm, a 6-fold increase (**Figure 4D**, **4D’**). These results confirm Sec16A as a driver for Sec body formation.

To further verify this, we visualized Sec body formation using mNeonGreen (mNG) fluorescently tagged Sec16A. Again, Sec body formation occurs quickly in cells expressing mNG-Sec16A upon 30- and 60-min incubation in KRBm (**Figure 4E, E’**). Importantly, incubation of cells with RPMI for 1 h after 1 h KRB treatment causes Sec body disassembly, leading to a re-localization of mNG-Sec16A similar to control cells (**Figure 4E, E’**). This result indicates that tagged-Sec16A induces formation of reversible Sec bodies.

### Sec16B is also a component of Sec body formation but not a driver

Mammalian genomes contain two distinct genes encoding for Sec16 proteins (Sec16A and Sec16B) (Bhattacharyya and Glick, 2007). Sec16B is 50 % smaller than Sec16A but both proteins localize to ERES in mammalian cells. Unlike Sec16A, Sec16B does not contain the SRDC in its C-terminus or anywhere else in its sequence, but it does display a similar extent of disordered regions, especially at the C-terminal region (Figure 5—figure supplement 1A). Interestingly, these two proteins have overlapping functions, yet do they not compensate for each other (Budnik et al., 2011).

In this respect, we questioned the role of Sec16B in Sec body formation. To do so, we specifically depleted it using specific siRNAs, leading to an 81 % reduction in *Sec16B* mRNA level (**Figure 5A**). However, in stark contrast with Sec16A depletion, Sec16B depletion does not prevent Sec body formation neither upon KRBm incubation (**Figure 5B, B’**) nor upon NaCl stress (Figure 5—figure supplement 1B, B’).

**Figure 5.**
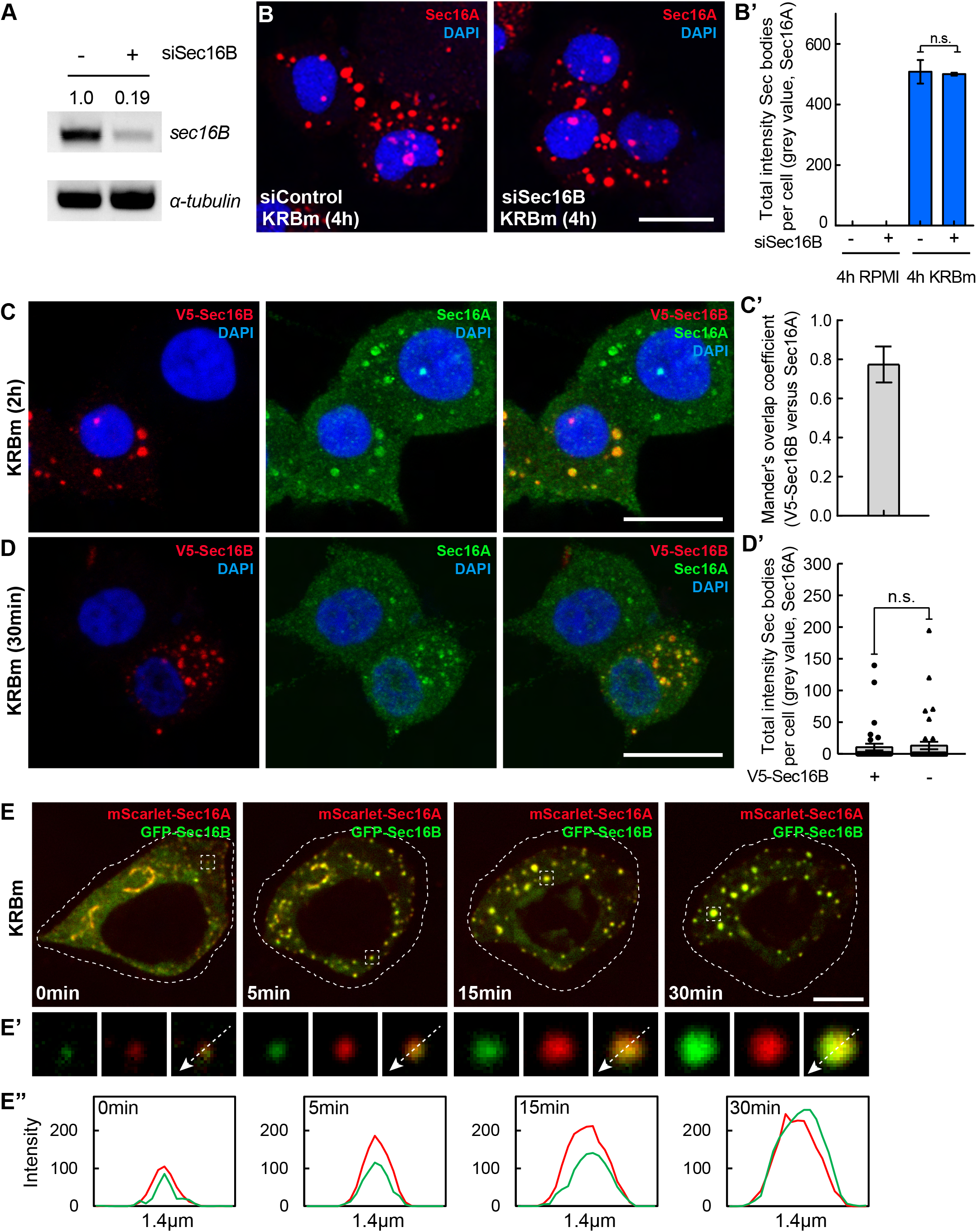
Sec16B is a component of Sec bodies but does not drive their formation. **A:** PCR of Sec16B and α-tubulin cDNAs from mock and Sec16B depleted INS-1 cells. Note that depleting Sec16B for 4 days leads to a very efficient knockdown (81%). Normalized value of Sec16B against α-tubulin, was used to calculate the Sec16B ratio between mock and Sec16B-depleted cells. **B, B’:** Immunofluorescence visualization of Sec16A in Sec16B depleted INS-1 cells incubated in KRBm for 4 h (B). Note that Sec body formation (marked by Sec16A) is not inhibited upon Sec16B depletion. Quantification of the total intensity of Sec bodies per cell (B’); n=2 experiments, both exp > 30 cells **C, C’:** Immunofluorescence visualization of V5-Sec16B (anti V5) and endogenous Sec16A in V5-Sec16A transfected INS-1 cell after KRBm incubation for 2 h (C). Mander’s overlap coefficient of V5 and Sec16A in Sec bodies (C’); n=51 cells. **D, D’:** Immunofluorescence visualization of V5-Sec16B (anti V5) and endogenous Sec16A in V5-Sec16B transfected INS-1 cell after KRBm incubation 30 min. Quantification of Sec16A-positive Sec body total intensity per cell in transfected and non-transfected cells in (D’); n=35 and 42 cells. **E-E”:** Representative still images of time-points 0, 5, 15 and 30 min from a live cell expressing mScarlet-Sec16A and GFP-Sec16B and treated with KRBm during live cell imaging (E). Single and merged channels of selected ERES and Sec body structures (E’). Intensity profile lines displaying co-distribution of Sec16A and Sec16B (E”). See also Video S2. See also Figure 5—figure supplement 1. Scale bar: 10 μm (C, D); 5 μm (E) Error bar: SD (C’), SEM (B’) (D’); ns – not significant

This result might result from the fact that Sec16B is not a Sec body component. To test this, we overexpressed V5-tagged Sec16B, which in control conditions localized to ERES, as expected (Figure 5—figure supplement 1C). Similar to Sec16A, Sec16B is efficiently incorporated into Sec bodies upon KRBm incubation for 2 h (**Figure 5C**) where it co-localizes with endogenous Sec16A **(Figure 5C**), with a Mander’s coefficient of around 0.77 (**Figure 5C’**). These results show that Sec16B is also a component of Sec bodies. However, unlike Sec16A (**Figure 4D, D’**), overexpression of V5-Sec16B does not promote Sec bodies formation after KRBm incubation for 30 min (**Figure 5D**, **5D’**). This suggests that Sec16B is a component of Sec bodies but not a driver of their formation.

Given this differential role in driving Sec body formation, we asked whether Sec16A and Sec16B are recruited to the same forming Sec bodies. To address this, we expressed fluorescently tagged mScarrlet-Sec16A and GFP-Sec16B and visualized transfected cells by live-cell imaging upon KRBm incubation. Critically, each Sec body contains both Sec16 species that perfectly co-localize and are recruited to Sec bodies with same kinetics (**Figure 5E-E’’**).

Taken together, these results strongly indicate that both Sec16A and Sec16B are components of Sec bodies, but that only Sec16A is a driver for stress-induced Sec body formation in mammalian cells.

### ERES components are simultaneously recruited into Sec bodies that form by fusion

We then questioned the mechanism behind the formation of Sec bodies in INS-1 cells. INS-1 cell Sec bodies are sensitive to hexanediol, suggesting that they are liquid droplets. One of the properties of liquid droplets is their propensity to fuse with one another, especially the small ones fusing with larger ones, the so-called ‘’droplet fusion’’ effect (Brangwynne et al., 2009; Weber and Brangwynne, 2012), leading to coalescence of a given size.

To study the dynamics of Sec body formation, we performed live-cell imaging of mNG-Sec16A every 30 seconds for 30 min during KRBm treatment. Live-cell imaging revealed that ERES started remodeling into small round foci within few minutes of KRBm treatment. These foci then fused with each other over time leading to the formation of typical Sec bodies within 30 minutes of treatment (**Figure 6A,** Video S1). We quantified the total number of fusion events occurring during 5 min, in intervals, until 30-min endpoint. Interestingly, we observed that the number of fusion events peaks at 10-15 min. After 20 min, fusion events are still observed but fewer as newly formed Sec bodies reach a stable size of 0.5-1 μm in diameter (**Figure 6B,** Video S1).

**Figure 6.**
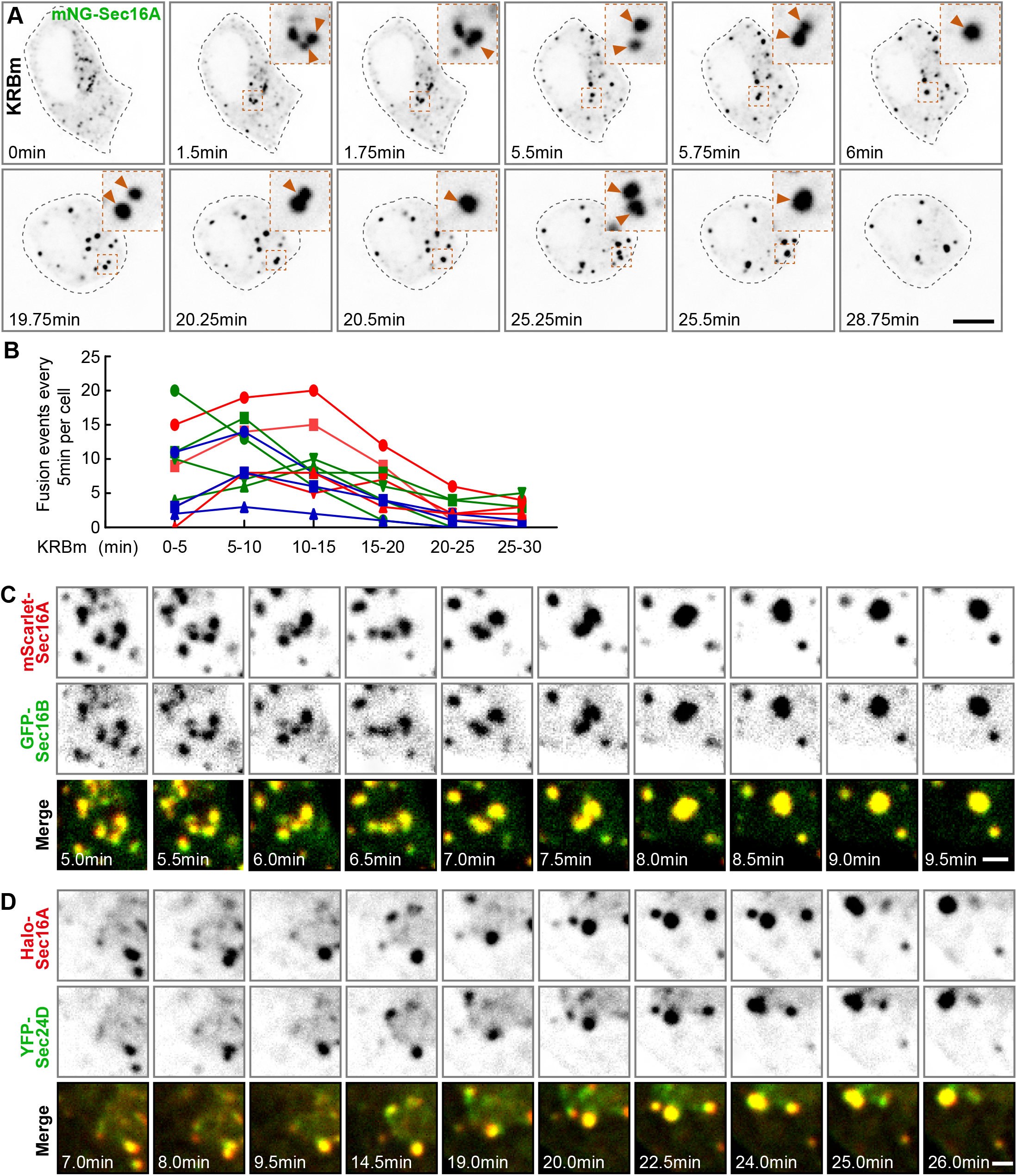
Sec body assembly is mediated by fusion of small foci containing several ERES components. **A**: Representative still images of a live INS-1 cell transfected with mNeonGreen (mNG)-Sec16A and recorded every 15 seconds for 30 min during KRBm incubation. Arrows in inserts despite fusion events. See also Video S1. **B**: Quantification of number of fusion events occurring during 5 min for 30min KRB incubation. In the graph, lines correspond to individual cells undergoing fusion upon KRBm treatment; n=11 cells. **C**: Representative still images of a region of a live INS-1 cell expressing mScarlet-Sec16A and GFP-Sec16B imaged every 30 seconds for 30 min during KRBm incubation. Note that mScarlet-Sec16A and GFP-Sec16B are simultaneously recruited into newly form Sec bodies. See also Figure 5E and Video S2. **D:** Representative still images of a region of an INS-1 cell expressing Halo-Sec16A and YFP-Sec24D. Cells were pre-incubated with the permeable Halo-646 dye prior to imaging. Live-imaged as in (C). Simultaneous recruitment of Halo-Sec16A and YFP-Sec24D into assembling Sec bodies is shown. See also Figure 6—figure supplement 1 and Video S3. Scale bar 5 μm (A), 1 μm (B) and (C).

We have shown that Sec16A and Sec16B co-localize in large Sec bodies (**Figure 5C, E-E’’**). We have also shown that Sec bodies contain COPII subunits (**Figure 2**). Using live-cell imaging, we then asked whether these different ERES components are either sequentially or simultaneously recruited into forming Sec bodies. We observed that mScarlet-Sec16A and GFP-Sec16B co-distribute in most of the small foci that are observed just after the beginning of the KRBm treatment (**Figure 5E-E’’; Figure 6C,** Video S2). These dually positive small foci undergo further fusion events until Sec bodies reached their large typical size (**Figure 6C,** Video S2). Similar results were observed in cells expressing Halo-Sec16A and YFP-Sec24D (**Figure 6D**, Figure 6—figure supplement 1, Video S3).

These results indicate that Sec bodies form by fusion of remodeled ERES and that the different ERES components are recruited simultaneously into newly formed Sec bodies.

### Testing the relationship between Sec bodies formation and ER exit

Last, we questioned the role of stress on the inhibition of the early secretory pathway. Indeed, extracellular stress is known to influence protein transport (Farhan and Rabouille, 2011; van Leeuwen et al., 2018).

More specifically, given that Sec bodies contain both Sec16 orthologs and most of the COPII subunits, we questioned the role of Sec bodies with regards to the inhibition of protein transport out of the ER. In *Drosophila* cells, the formation of Sec bodies in stressed cells correlates with transport inhibition in the secretory pathway (Zacharogianni et al., 2014). However, the questions as to whether Sec body formation causes inhibition of protein transport (and ER exit in particular) and whether Sec bodies could act as transport intermediates remain to be answered.

To test whether Sec body formation in INS-1 cells coincides/correlates with inhibition of trafficking in the early secretory pathway, we used the RUSH system to study the ER exit of the Transferrin receptor (TfR) (Boncompain et al., 2012). As expected, Strep-KDEL-Halo-Transferrin Receptor (TfR)-SBP (RUSH-TfR) is retained in the ER when cells are incubated in DMEM without biotin (Figure 7—figure supplement 1A). The addition of biotin for 30 minutes leads to the release of RUSH-TfR out of the ER in 93 % of the cells (**Figure 7A, 7A’**) and reaches a compartment largely positive for Giantin (Figure 7—figure supplement 1B).

**Figure 7:**
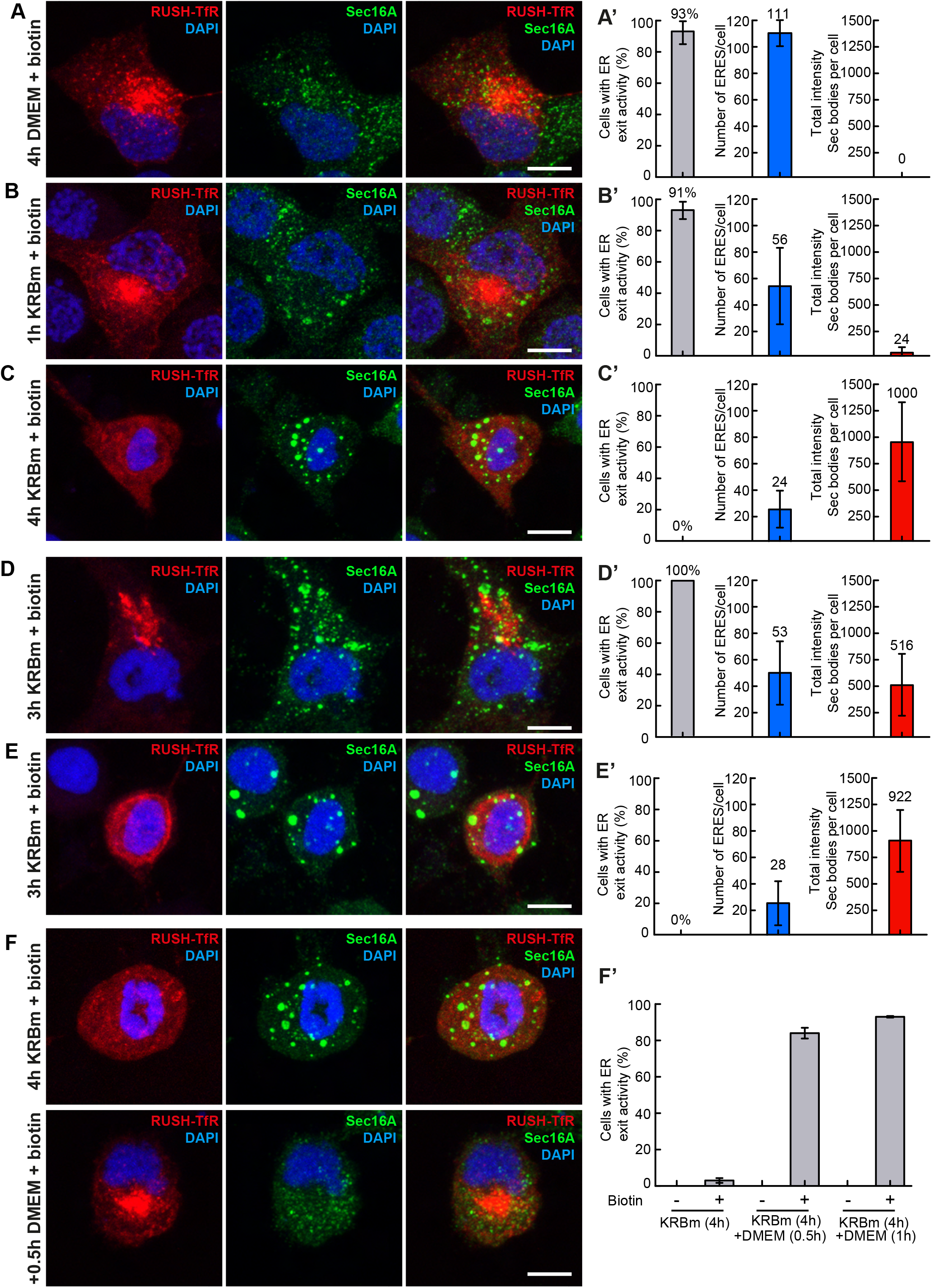
Sec body formation precedes ER exit. **A, A’**: Visualization of RUSH-TfR and endogenous Sec16A upon addition of biotin (30 min, 100 μM) in DMEM (4 h) cultured INS-1 cells. Quantification of the ER exit activity, number ERES and Sec bodies (A’); n=16 cells. **B, B’**: Visualization of RUSH-TfR and endogenous Sec16A upon addition of biotin (30 min, 100 μM) in INS-1 cells in KRBm for 1 h. Quantification of the ER exit activity, number ERES and Sec bodies (B’); n=30 cells. **C, C’**: Visualization of RUSH-TfR and endogenous Sec16A upon addition of biotin (30 min, 100 μM) in cells in KRBm for 4 h. Quantification of the ER exit activity, number ERES and Sec bodies (C’); n=12 cells. **D-E’**: Visualization of RUSH-TfR and endogenous Sec16A upon addition of biotin (30 min, 100 μM) in cells in KRBm for 3 h. The release of RUSH-TfR out of the ER is observed in 50% of the cells (D, D’), and RUSH-TfR retention in the ER is observed in the remaining cells (E, E’). Quantification of the ER exit activity, number ERES and Sec bodies (D’, E’); n=27 cells. **F, F’**: Visualization of RUSH-TfR and endogenous Sec16A in cells incubated in KRBm for 4 h followed by 30 min (0.5 h) or 1 h in DMEM (including 30 min biotin, 100 μM). Note that the Sec bodies have dissolved and that RUSH-TfR has efficiently exited the ER. Quantification of the ER exit (F’); n=2 experiments. See also Figure 7—figure supplement 1 Scale bar: 10 μm (A-F) Error bar: SD (A’-E’), SEM (F’)

We then asked what happens to protein exit from the ER in cells upon amino acid starvation in KRBm. To assess this, we incubated transfected cells for increasing times in KRBm and then added biotin for the last 30 min of this incubation. If ER exit is inhibited upon stress, RUSH-TfR is retained in the ER, even after the addition of biotin. If the ER exit is still active, it will exit the ER.

After 1 h KRBm (including 30 min biotin), RUSH-TfR exits the ER and reaches the Golgi in 91 % of the cells, showing that ER exit is still active (**Figure 7B, B’**). In the absence of biotin, RUSH-TfR is retained in the ER as expected (Figure 7—figure supplement 1C). In this population of cells, hardly any Sec bodies have formed (**Figure 7B, B’,** red bar). This shows that the mere incubation in KRBm is not sufficient to impede ER exit. In contrast, after 4 h in KRBm (including 30 min biotin), RUSH-TfR is retained in the ER in 100 % of the cells, showing that ER exit is fully inhibited (**Figure 7C, C’**) and these cells exhibit the full complement of Sec bodies (**Figure 7C, C’,** red bar). Therefore, as in *Drosophila* S2 cells (Zacharogianni et al., 2014), these results show a strong correlation between the formation of Sec bodies and inhibition of ER exit. Furthermore, as we have not observed a co-localization of RUSH-TfR to Sec bodies, these results also demonstrate that Sec bodies are transport incompetent.

### Sec body formation control ER exit by regulating the number of functional ERES

To clarify whether ER exit inhibition is a consequence of Sec body formation or the other way around, i.e., whether Sec body formation is a consequence of ER exit blockade, we examined the cells after 3 h incubation in KRBm (including 30 min biotin). There, RUSH-TfR appears to exit the ER in 49 % of the cells, with a moderate Sec body formation (**Figure 7D, D’**). In the other 51 % of the cells, RUSH-TfR is retained in the ER and Sec body have formed as efficiently as in 4 h of KRBm (**Figure 7E, E’**). Taken together, these results indicate that Sec body formation is not a consequence of the inhibition of ER exit. Intriguingly, the number of Sec bodies does not seem to be a key factor in ER exit, as both populations of cells in 3 h KRBm (with and without ER exit activity) form Sec bodies (6.5 and 7.4, respectively; **Figure 7D’, E’**).

To further understand what distinguishes these two populations of cells, these where ER exit occurs from those where it does not, we quantified the number of ERES in cells of each category. In cells treated with KRBm for 3 h where RUSH-TfR exits the ER, the number of ERES is half of what it is in DMEM (53 versus 111/cell, respectively) (**Figure 7A’, D’,** blue bars), a number that is comparable to the cells incubated in KRBm for 1 h (56), where ER exit is very efficient (compare blue bars in **Figure 7B’ and D’**). The decrease in the ERES number from 111 to 53 upon 1 h KRBm, is likely due to their initial remodeling into slightly larger and brighter structures (Figure 7—figure supplement 1D) and may correspond to the first fusion events observed by live-cell imaging (**Figure 6A).** In contrast, in cells where ER exit does not occur, the number of ERES has dropped to 28 (**Figure 7E’**), a number very similar to cells incubated in KRBm for 4 h (24) where ER exit is fully blocked (**Figure 7C’**).

These results show that Sec bodies formation precedes the inhibition of ER exit and show that Sec body formation is not a consequence of ER exit inhibition. Instead, Sec body formation appears to cause the inhibition of ER exit. We propose that cells are competent for ER exit until the number of ERES has dropped by 75% of what it is in DMEM. This suggests that Sec body formation recruits Sec16A (and the other COPII subunits) away from ERES, leading to their progressive depletion from ERES that become non-functional. In other words, ER exit is inhibited when Sec16A and COPII subunits are significantly incorporated into Sec bodies and no longer available at ERES.

To test this further, we monitored RUSH-TfR transport upon stress relief (i.e., in cells incubated in KRBm for 4 h followed by either 30 min or 1 h back in DMEM). This condition leads to the complete dissolution of Sec bodies and the re-localization of Sec16A to ERES (**Figure 3B, Figure 7F**). Strikingly, this is accompanied by the exit of RUSH-TfR from the ER in 90% of the cells upon biotin addition (**Figure 7F’**) whereas it is properly retained in the ER when biotin is not added (Figure 7— figure supplement 1E).

Taken together, this experiment shows that in INS-1 cells, Sec bodies act as a storage for critical ERES components, such as Sec16A, controlling its availability to ERES functioning in ER exit. It also shows that ERES components are quickly available again upon stress relief, leading to active protein transport.

## Discussion

Remodeling of ERES into membraneless Sec bodies have been observed in *Drosophila* S2 cells upon stress. In this study we report that Sec bodies also form in mammalian cells upon same stress-specific conditions. Mammalian Sec bodies share several characteristics with *Drosophila* Sec bodies. For instance, they recruit Sec16 protein(s) and COPII subunits, are membranelles and reversible structures, which indicate they are evolutionary conserved. In mammalian cells, the two Sec16 ortholog, Sec16A and 16B, are both recruited into Sec bodies but only Sec16A is a Sec body driver. The remodeling of ERES into Sec bodies occurs by fusion events, in which ERES components are simultaneously recruited into forming Sec bodies. Lastly, we propose that the remodeling of ERES into Sec bodies causes a depletion of functional ERES, thus shutting down the early secretory pathway during stress, which is efficiently restored after stress relief. Given the relationship between stress assemblies and the modulation of anabolic pathways, it is likely that Sec body formation is a key conserved mechanism of cell survival during stress and fitness upon stress relief (Kroschwald and Alberti, 2017; van Leeuwen and Rabouille, 2019) through their capacity of modulating protein secretion.

### Sec bodies are conserved in evolution. A key stress response?

In contrast to stress granules that form upon different types of stress (heat shock, ER stress, sodium arsenite, osmotic stress) (Aulas et al., 2018; van Leeuwen and Rabouille, 2019), studies in *Drosophila* cells have shown that Sec bodies form only upon specific types of stress, i.e. high NaCl stress and upon amino acid starvation in KRB (Zhang et al., 2021). In mammalian INS-1 cells the same stresses also induce a strong remodeling of ERES into Sec bodies that have very similar features as their invertebrate counterparts, revealing a strong convergent evolution potentially underlying a fundamental ancient stress response. Interestingly, Sec16 and all COPII subunits are conserved in last eukaryotic common ancestor, suggesting that they could also respond to stress in ancient cellular lineages (Schlacht and Dacks, 2015).

It remains unclear whether Sec bodies can form upon different types of stress in different mammalian cells and why only a modest remodeling of ERES in HepG2, MDCKII and MRC5 cells was observed upon high salt stress whereas INS-1 cells respond strongly. At present, the physiology behind these differences is not clear. It could be related to the secretory capacities of these cells, or the amount of the Sec body driver, Sec16A (the isoform containing the SRDC domain).

### Sec body form by fusion in close proximity to the ER

One important feature of membraneless organelles is that they are not enclosed by a lipid membrane. Sec bodies are not delimited by membrane, but they are in close proximity to the ER membrane. This interaction with the ER is consistent with the first step of their biogenesis through the fusion of ERES. This suggests that the ERES components that form Sec bodies do not disperse prior to their coalescence but rather coalesce in situ. The remodeled small structures (larger ERES) continue to fuse each other to form small and then large Sec bodies, until they reach an optimal and stable size around 1 μm in diameter. Another characteristic of membraneless organelles is that they are reversible. We observed that Sec bodies efficiently disassemble and restore ERES, even after just 30 min of stress removal. This fast and efficient response to stress-relief may be related to the close contact with the ER membrane. It remains elusive how Sec bodies disassemble after stress relief, either through dispersion of ERES components, perhaps by active signaling as for stress granules (Wippich et al., 2013) or through fission events. Supporting the last model, recent findings have identified a role of the ER in membraneless P-bodies size by promoting fission events (Lee et al., 2020). It is likely that the close contact of Sec bodies with the ER facilitates a fast restoration of ERES on ER membrane.

### Sec bodies do not contain Golgi proteins

We have observed different ERES components recruited into INS-1 Sec bodies, such as Sec16A, Sec16B and the COPII subunits Sec13 and Sec24D (and likely Sec31 as in *Drosophila* S2 cells, (Zacharogianni et al., 2014)). Here, we show that ERES components are simultaneously recruited to forming Sec bodies, in agreement with the initial coalescence of functional ERES containing all required components. Conversely, Golgi proteins are largely excluded from Sec bodies, except for a small fraction of p115. p115 is associated to the ER-to-Golgi transport machinery and has been shown to regulate ERES (Alvarez et al., 1999; Alvarez et al., 2001; Kondylis and Rabouille, 2003; Sapperstein et al., 1995). It is therefore likely that the ERES-located p115 is incorporated into Sec bodies. This is also consistent with a recent proteomic analysis of Sec bodies in *Drosophila* S2 cells, where p115 is also found as part of the Sec body proteome (Zhang et al, 2022, submitted). Interestingly, GRASP55 and 65 appear to form larger structures upon stress but those are not Sec bodies (marked by Sec16). Whether those are related to CUPS (Compartments for Unconventional Protein Secretion), which also form upon nutrient starvation remains to be elucidated (Cruz-Garcia et al., 2014). This suggests that nutrient stress leads to an important remodeling of the cytoplasm that somehow remain distinct and sustain different functions.

### Sec body formation is driven by Sec16A

Phase-separation is often promoted by driver proteins. These proteins induce phase-separation through a mild change in their conformation. This leads to their coalescence that also recruits other proteins through low affinity interaction (Banani et al., 2017). The absence of drivers prevents phase separation (Banani et al., 2017; Banani et al., 2016).

Here we show that although the two mammalian Sec16 ortholog are present in INS-1 cells, Sec16A but not Sec16B is a driver of Sec bodies. In this regard, we have previously shown that *Drosophila* Sec16 (dSec16) is a driver in Sec body formation in *Drosophila* cells (Aguilera-Gomez et al., 2016), and mammalian Sec16A shares a large homology with *Drosophila* Sec16 (dSec16) (Ivan et al., 2008). As dSec16, Sec16A contains low complexity intrinsically disordered sequences that could potentially stimulate Sec body formation by engaging low affinity multivalent interactions with other ERES proteins. However, Sec16B also contains such disordered sequences, yet it is not a driver. This suggests that the presence of disordered sequences in Sec16A and Sec16B are likely necessary for their incorporation into Sec bodies, but not sufficient for driving their coalescence. Instead, we have shown that a conserved 44 residue domain called SRDC that is present in the C-terminus of dSec16 is instrumental for Sec body formation in S2 cells. When expressed on its own stimulate Sec body formation even in the absence of stress. Conversely, overexpression of dSec16 lacking a SRDC domain prevents Sec body formation (Aguilera-Gomez et al., 2016). Critically, Sec16A, but not Sec16B, also contains the SRDC, and Sec16A overexpression accelerates Sec body formation. We propose that the presence of the SRDC in Sec16A is potentially the driving factor in Sec body formation in INS-1 cells. Whether other Sec body components contribute to Sec body formation remains to be investigated.

### Sec bodies and ER exit

Here, we find that Sec body formation precedes inhibition of ER exit and protein secretion in the secretory pathway, showing that they are not the consequence of this ER inhibition. In other words, ER exit persists even when cells have already formed Sec bodies, as long as there are enough functional ERES. Instead, it appears that ER exit is inhibited when the number of functional ERES drops below a threshold and when their components (such as Sec16A and COPII subunits) are quantitatively recruited to Sec bodies. Sec16A is critical in COPII dynamics (Bhattacharyya and Glick, 2007; Tillmann et al., 2015; Hughes et al., 2009; Joo et al., 2016; Wilhelmi et al., 2016). Without Sec16A and COPII components at ERES, protein transport is slowed down, and cells proliferation is compromised (Tillmann et al., 2015).

This raises the question why Sec bodies form (recruiting both Sec16 and COPII subunits). From studies in *Drosophila* S2 cells, we have previously proposed that proteins recruitment into Sec bodies protect them from stress-mediated degradation in such a way that they are available upon stress relief. Here, we show that Sec body formation shut down ER exit. As secretion is an energy consuming process (Brandizzi and Barlowe, 2013; Depaoli et al., 2019; Sudhof and Rothman, 2009; Yong et al., 2019), the inhibition of its first step through Sec body formation would save energy that could be redirected to other processes important to cope with the stress itself. This is in line with the general notion that cellular stress leads to a slowing down of secretion (Farhan and Rabouille, 2011; Tillmann et al., 2013).

### Model: Sec bodies – ER exit – survival

We propose that Sec bodies also store key ERES components in a near native state, allowing them to be quickly functional upon stress relief. As a driver, Sec16A is likely slightly modified upon stress, perhaps through stress-induced reversible post-translational modifications, such as MARylation (Aguilera-Gomez et al., 2016). This would lead to a slight change of its conformation, resulting in the initiation of its coalescence that would promote the recruitment of the nearby ERES components, including COPII subunits with which it interacts (reviewed in (Sprangers and Rabouille, 2015)). This would deplete ER membrane from functional ERES, reducing the efficiency of protein transport out of the ER. The steps involved in stress-induced assembly of Sec bodies and ER exit inhibition is quickly and efficiently reverted upon stress removal, suggesting that the stress-induced Sec16A modifications are reversible, and that Sec bodies act as a storage for functional near native proteins. Together these data show that protein secretion inhibition is controlled by the dynamic interconversion of ERES and Sec bodies during periods of stress and stress relief.

This reinforces the notion that Sec bodies are pro-survival (as shown in *Drosophila* cells) in line with the general concept that the formation of stress assemblies is a key cellular mechanism to cope with stress and thrive upon stress relief (Kroschwald and Alberti, 2017; Riback et al., 2017).

## Material & Methods

### Cell culture and treatments

INS-1 cells and INS-1 823/3 cells (Sigma-Aldrich, scc208) were cultured in RPMI-1640 medium, GlutaMAX Supplement, HEPES (Gibco, 72400047), 10% FBS (Sigma-Aldrich, F7524), 1% penicillin/streptomycin (Gibco, 15140122) and 50 μM 2-Mercaptoethanol (Sigma-Aldrich, M6250). Cell were maintained at 37°C and 5% CO_2_.

Sec bodies were formed in INS-1 cells by incubating cells with either RPMI-1640 medium plus 200 mM NaCl (RPMI200) or Krebs Ringer Bicarbonate buffer modified (KRBm) for 4 hours. KRBm contains 0.7 mM Na_2_HPO_4_, 1.5 mM NaH_2_PO_4_, 30 mM NaHCO_3_, 237 mM NaCl, 4.53 mM KCl, 0.53 mM MgCl and 11 mM D-glucose at pH 7.4 (adjusted with HCl).

For overexpression of plasmids containing mNeongreen-Sec16A, mScarlet-Sec16A, GFP-Sec16B, Halo-Sec16A and YFP-Sec24D, INS-1 cells were transfected with Lipofectamine 2000 (Invitrogen, 11668019). DNA and Lipofectamine were mixed in OptiMEM (Gibco, 31985047) before being added to the cells. Cells were analyzed after 24-48h of transfection.

For knockdown experiments, INS-1 cells were transfected in 6-well plates with 5 nM siRNA using Lipofectamine RNAiMAX (Invitrogen, 13778030) and typically analyzed after 96 hours. The siRNAs used were directed against Sec16A (rat) (Ambion, s137444) and Sec16B (rat) (Ambion, s137156).

### DNA constructs

The following vectors were used: pCMV-mScarleti-C1 was provided by Dr. Dorus Gadella (Bindels et al., 2017) (Addgene plasmid #85044). pCMV-EGFP-C1 was a gift from Dr. Jennifer Lippincott-Schwartz. pCMV-EYFP-Sec24 (Stephens et al., 2000) (Addgene plasmid #66614), pCMV-EGFP-Sec16B (Budnik et al., 2011) (Addgene plasmid #66607) were provided by Dr. David Stephens. Strep-KDEL-Halo-Transferrin Receptor (TfR)-SBP (Halo-RUSH-TfR) (called RUSH-TfR in the text) was provided by Dr. Jennifer Lippincott-Schwartz (Weigel et al., 2021) (Addgene plasmid #166905) pF282-hEF1a-H2B-mNeonGreen-IRES-Puro x Tol2 was a gift from Dr. Judith Klumperman. Halo-Clathrin (Catsburg et al., 2022) was a gift from Dr. Harold MacGillavry.

The following plasmids were generated in this study by Gibson assembly, using NEBuilder® HiFi DNA Assembly Master Mix (NEB, E2621L). To obtain pCMV-mNeonGreen-Sec16A, we first generated the pCMV-EGFP-Sec16A construct. In detail, rat Sec16A (NM_001276417.1) was PCR amplified from a cDNA library obtained from INS-1-derived mRNA and inserted into pCMV-EGFP-C1 vector between XhoI and KpI restriction sites. After generating pCMV-EGFP-Sec16A, EGFP sequence from pCMV-EGFP-Sec16A was removed by restriction digest with AgeI and BsrGI and replaced with a mNeonGreen sequence. The mNeonGreen sequence was PCR amplified from the template hEF1a-H2B-mNeonGreen-IRES-Puro x Tol2, and a Kozak sequence was also added in front of mNeonGreen to increase the efficiency of translation. A 6-amino acid flexible linker (SGLRSR) was introduced between mNeonGreen and the beginning of Sec16A by adding additional nucleotides to the cloning primers.

For pCMV-Halo-Sec16A, the construct was generated in a similar way as described above for pCMV-mNeonGreen-Sec16A. In detail, the EGFP sequence from pCMV-EGFP-Sec16A was removed by a digestion with AgeI and BsrGI and replaced with a Halo tag sequence. The Halo tag sequence was amplified by PCR from the template pHalo-Clathrin, and a Kozak sequence was added in front of the Halo tag sequence. A flexible linker of 7 amino acids (KSGLRSR) was flanked between Halo tag sequence and the beginning of Sec16A by including extra nucleotides to the cloning primers.

For pCMV-mScarlet-Sec16A, rat Sec16A sequence was amplified by PCR from pCMV-EGFP-Sec16A and inserted into pCMV-mScarleti-C1 vector between BamHI and BglII sites. A flexible linker of 6 amino acids (SGLSGS) was introduced between the mScarleti sequence and the beginning of Sec16A sequence by adding extra nucleotides to the cloning primers.

For pCMV-V5-Sec16A, a V5 sequence was amplified and inserted into pCMV-EGFP-Sec16A cut open with AgeI and BsrGI to replace the GFP sequence. A Kozak sequence and a flexible linker (GPKSGLRSR) were added in front and behind the V5 sequence, respectively.

For pCMV-V5-Sec16B, similarly, a V5 sequence was amplified and inserted into pCMV-BFP-Sec16B (generated in our lab) cut open with AgeI and HindIII. A Kozak sequence was added in front of V5, and a flexible linker (GPTNGSGSGS) was introduced behind V5 sequence. All primers used in this study are listed in *Supplemental Table S1*.

### Antibodies

For immunofluorescence, we used the primary antibodies rabbit anti-Sec16A (1:400) (Bethyl, A300-648A), mouse anti-Sec13 (1:100) (Santa Cruz, sc-514308), mouse anti-Sec24A (1:100) (Santa Cruz, sc-517155), mouse anti-p115 (1:100) (gift from Martin Lowe, University of Manchester, UK), mouse anti-GM130 (1:500) (gift from Martin Lowe, University of Manchester, UK), rabbit anti-GRASP55 (1:1000) (Gift from M. Bekier, University of Michigan, MI, USA) (Xiang and Wang, 2010), rabbit anti-GRASP65 (1:100) (Gift from F.A. Barr, Mac Planck Insititut für Biochemie, Planegg, Germany) (Shorter et al., 1999), mouse-anti V5 (Invitrogen, R96025). Donkey anti-rabbit Alexa 568 (1:500) (Invitrogen, A10042), goat anti-mouse Alexa 488 (1:500) (Invitrogen, A11001), donkey anti-mouse Alexa 555 (Invitrogen, A31570), donkey anti-rabbit (Invitrogen, A21206), goat anti-mouse IgG2a (Invitrogen, A21131), and goat anti-mouse IgG1 (Invitrogen, A21125) were used as secondary antibodies.

### Immunofluorescence

Cells were fixed in 4% paraformaldehyde in PBS (pH 7.4) for 15 minutes at room temperature. Then, cells were washed 3 times in PBS with 20 mM glycine (PBS-G) followed by permeabilization in PBS-G with 0.1% Triton X-100 for 10 minutes at room temperature. Subsequently, cells were washed 3 times with PBS-G and blocked in PBS-G with 0.5% fish skin gelatin (Sigma-Aldrich, G7041) for 20 minutes at room temperature. Next, cells were incubated with the primary antibody (in blocking buffer) for 1 hour at room temperature, followed by 3 times washing with blocking buffer. Then, cells were incubated with the secondary antibody for 1 hour at room temperature in the dark. Finally, cells were washed 3 times with PBS and coverslips were mounted on a microscope slide with Prolong Gold Antifade reagent with DAPI (Invitrogen, P36935).

### Microscopy and image acquisition

Fixed INS-1 cells were imaged on the laser scanning confocal microscope Leica SP8 or Zeiss LSM700. All images were acquired using a 63x oil immersion objective (NA 1.4).

For live-cell imaging experiments, we used an inverted microscope Nikon Eclipse Ti-E (Nikon), equipped with a Plan Apo VC ×100 NA 1.40 oil objective (Nikon), a Yokogawa CSU-X1-A1 spinning disk confocal unit (Roper Scientific), a Photometrics Evolve 512 EMCCD camera (Roper Scientific) or Photometrics Prime BSI camera, and an incubation chamber (Tokai Hit) mounted on a motorized XYZ stage (Applied Scientific Instrumentation). To control all devices, MetaMorph (Molecular Devices) version 7.10.2.240 software was installed. For Halo-tag, cells were pre-incubated with Janelia Fluor® 646 HaloTag® Ligand (100 nM) for 30 min, followed by a washing in RPMI-1640 medium prior to imaging. Coverslips were then mounted in a metal Ludin Chamber-Type I and supplemented in the original medium from INS-1 cells were imaged in a Tokai Hit incubation chamber that maintains optimal temperature and CO2 (37 °C and 5% CO2). To visualize different fluorescently tagged-proteins a laser channel was exposed for 200–300 ms while for multi-color acquisition, different laser channels were exposed for 200–400 ms sequentially. Cells were then washed once and imaged in KRBm every 15 or 30 seconds for 30 minutes. Total time and intervals of imaging acquisition for each experiment are depicted in each legend for figure and/or legend for Videos

### Quantification of Sec16A remodeling (Sec body formation)

Sec16A remodeling (and further Sec body formation) in INS-1 cells were quantified by using ImageJ (Schneider et al., 2012). In brief, a maximum z-projection was obtained from single Z-plane images. Next, a threshold was set using the function ‘’set threshold’’ with various parameters depending on the experiment. Then the intensity and particle size were measured with the tool ‘’Analyze particles’’. The intensity and size of each particle from one condition were stacked in a large matrix and the total intensity per particle was calculated by multiplying the area*intensity. We defined that a “large” Sec body has a particle size of >0.3 μm and an intensity of > 80% of the max intense particle. A “small” Sec body was defined with a particle size between 0.15-0.3 μm and an intensity of > 75% of the max intense particle. The total intensity of all ‘’small’’ and ‘’large’’ Sec bodies were summed up together to get the total intensity of all Sec bodies. The total intensity of Sec bodies per cell was represented by: (1) dividing the total intensity from all Sec bodies (respective to their criteria) by the total amount of cells; or (2) mean values of total intensity of Sec bodies per cell from all cells.

For colocalization of two different proteins, Mander’s coefficient (ImageJ) was used. Profile lines plots were generated by tracing a line along a Sec body using the RGB Profile Plot plugin in ImageJ.

### RUSH assay to assess protein exit out of the ER

INS-1 cells (P60-P80) were conditioned in Dulbecco’s modified Eagle medium (DMEM) (Invitrogen), GlutaMAX Supplement, 10% FBS (Sigma-Aldrich, F7524) and 1% penicillin/streptomycin (Gibco, 15140122) for 3 days at 37 °C and 5% of CO_2_ because RPMI contains high amount of biotin preventing the retention of RUSH-TfR in the ER even in the absence of exogenous added biotin. Cells were plated to 50% confluency on coverslips in a 12-wells plate 1 day before transfection. Per well, INS-1 cells were transfected with RUSH-TfR and Lipofectamine 2000 in OptiMEM. Subsequently, cells were washed and incubated in DMEM for 24 h. Cells were treated as indicated in the results. Unhooking of RUSH-TfR from the ER was allowed by the addition of D-biotin (Sigma-Aldrich, B4501-500MG) at 100 μM final concentration in DMEM for 30 min prior fixation. RUSH-TfR was visualized by staining cells with Janelia FluorX® 554 HaloTag® Ligand (100nM) together with the secondary antibody to label other proteins in blocking buffer for 1h at RT.

### Statistical analysis

Data obtained at least from two independent experiments was processed and statistically analyzed using Excel and Graphpad Prism. Mann-Whitney and Kruskal-Wallis test followed by a Dunn’s multiple comparison test were performed for statistical analysis and are indicated in Figure legends. Significance was determined as followings: ns-not significant, *p<0.05 **p<0.01 ***p<0.001. The assumption of data normality was evaluated using D’Agostino-Pearson omnibus test.

## Acknowledgments

We thank Sergio Reijnders (Hubrecht Institute) for helping to establish the quantification of Sec body formation; Anko de Graaff from the Hubrecht Imaging Facility for helping with the imaging; and Chun Hei Li (Utrecht University) for helping and optimizing the RUSH system in INS-1 cells.

## Funding

This work was supported by the European Union’s Horizon 2020 research and innovation programme under the Marie Skłodowska-Curie grant agreement (ITN-SAND 860035) and European Research Council grant (ERC-StG 950617) to G.G.F.

---

**Figure 1—figure supplement 1:**
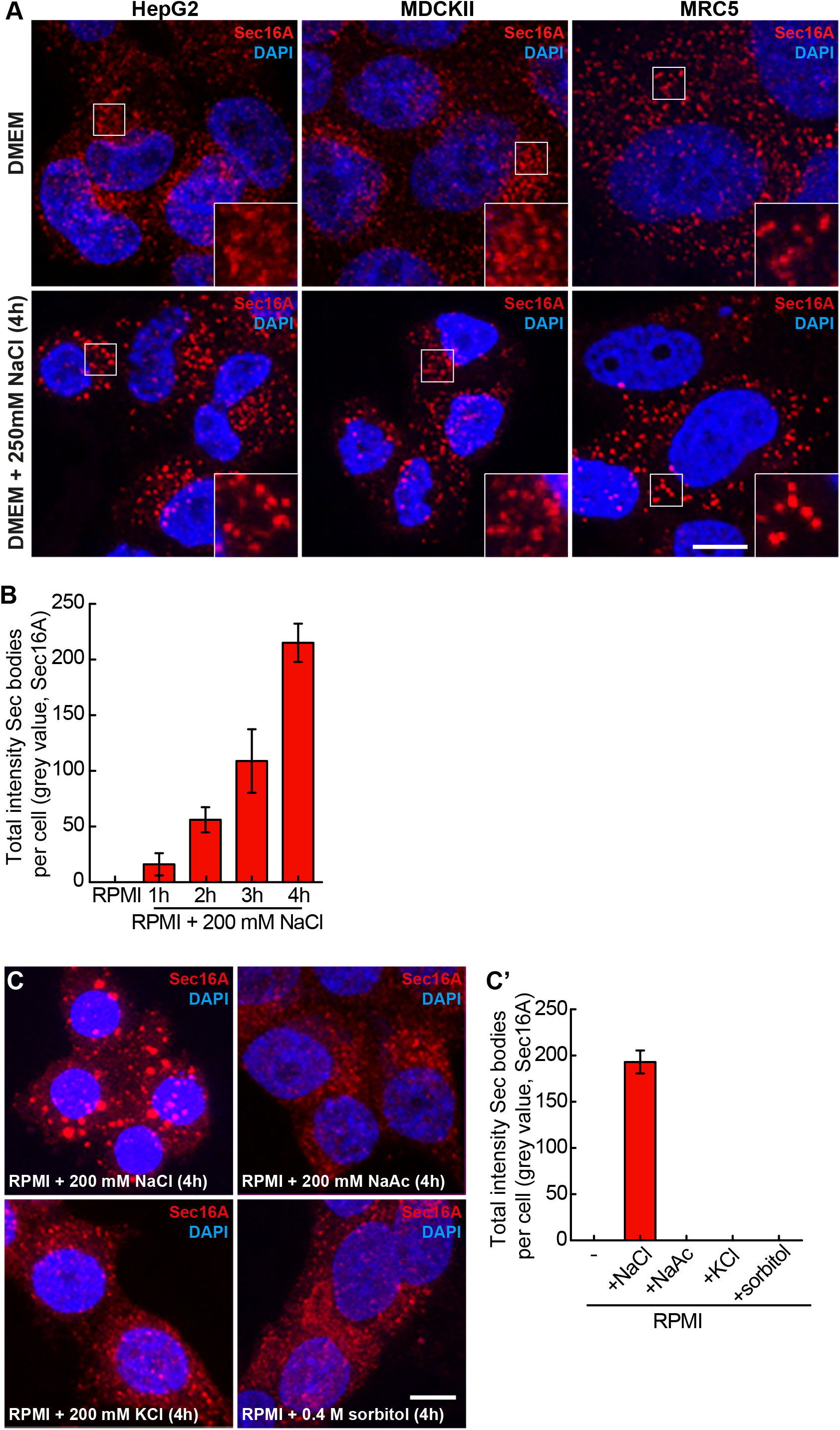
NaCl stress leads to mild remodeling of ERES in HepG2, MDCKII and MRC5 cells and strong remodeling in INS-1 cells. **A:** Immunofluorescence visualization of endogenous Sec16A in HepG2, MDCKII and MRC5 cells upon DMEM and DMEM+250 mM NaCl (4 h). **B:** Bar plot depicting the total intensity of Sec bodies per cell in RPMI200 over time (1-4 h) in INS-1 cells; n=2 experiments, both exp > 30 cells. **C, C’**: Immunofluorescence visualization of Sec16A in INS-1 cells upon incubation in RMPI supplemented with either NaCl, or Na-Acetate or KCl (200 mM, 4 h), or 0.4 M sorbitol for 4 h (C). Note that large Sec16A-positive structures only form upon the addition of NaCl. Quantification of the total intensity of Sec16A-positive remodeled structures per cell (C’); n=2 experiments, both exp > 30 cells. Scale bar: 10 μm (A) and (C). Error bar: SEM (B) and (C’)

**Figure 2—figure supplement 1:**
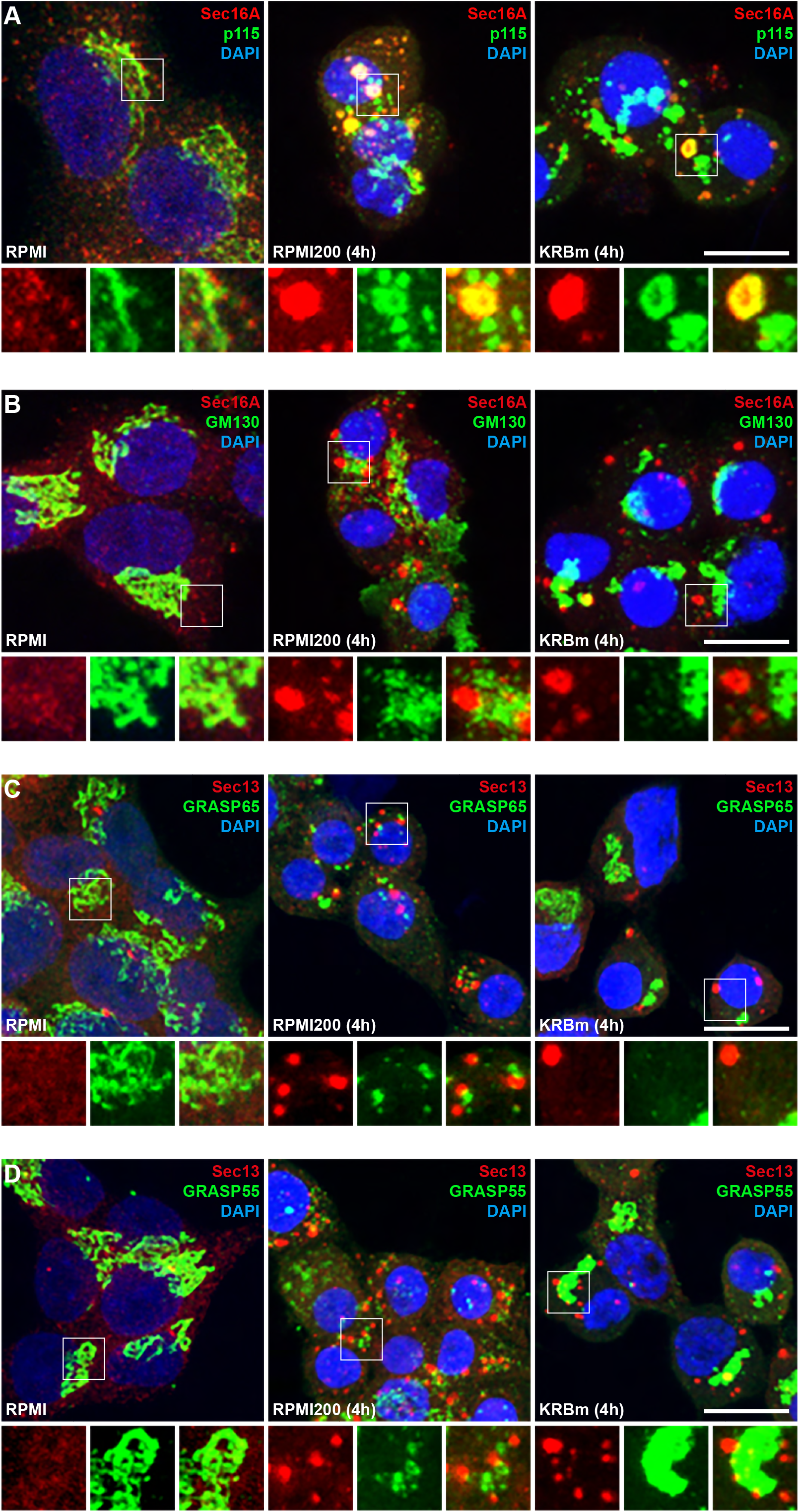
Sec16A-positive structures do not contain Golgi proteins. **A:** Immunofluorescence visualization of Sec16A and p115 in INS-1 cells upon incubation in RPMI, RPMI200 and KRBm (4 h). Note that a small fraction of p115 localizes in Sec16A positive structures. **B:** Immunofluorescence visualization of Sec16A and the Golgi marker GM130 in INS-1 cells upon incubation in RPMI, RPMI200 and KRBm (4 h). **C:** Immunofluorescence visualization of Sec13 and GRASP65 in INS-1 cells upon incubation in RPMI, RPMI200 and KRBm (4 h). **D:** Immunofluorescence visualization of Sec13 and GRASP55 in INS-1 cells upon incubation in RPMI, RPMI200 and KRBm (4 h). Scale bar: 10 μm (A-D)

**Figure 4—figure supplement 1.**
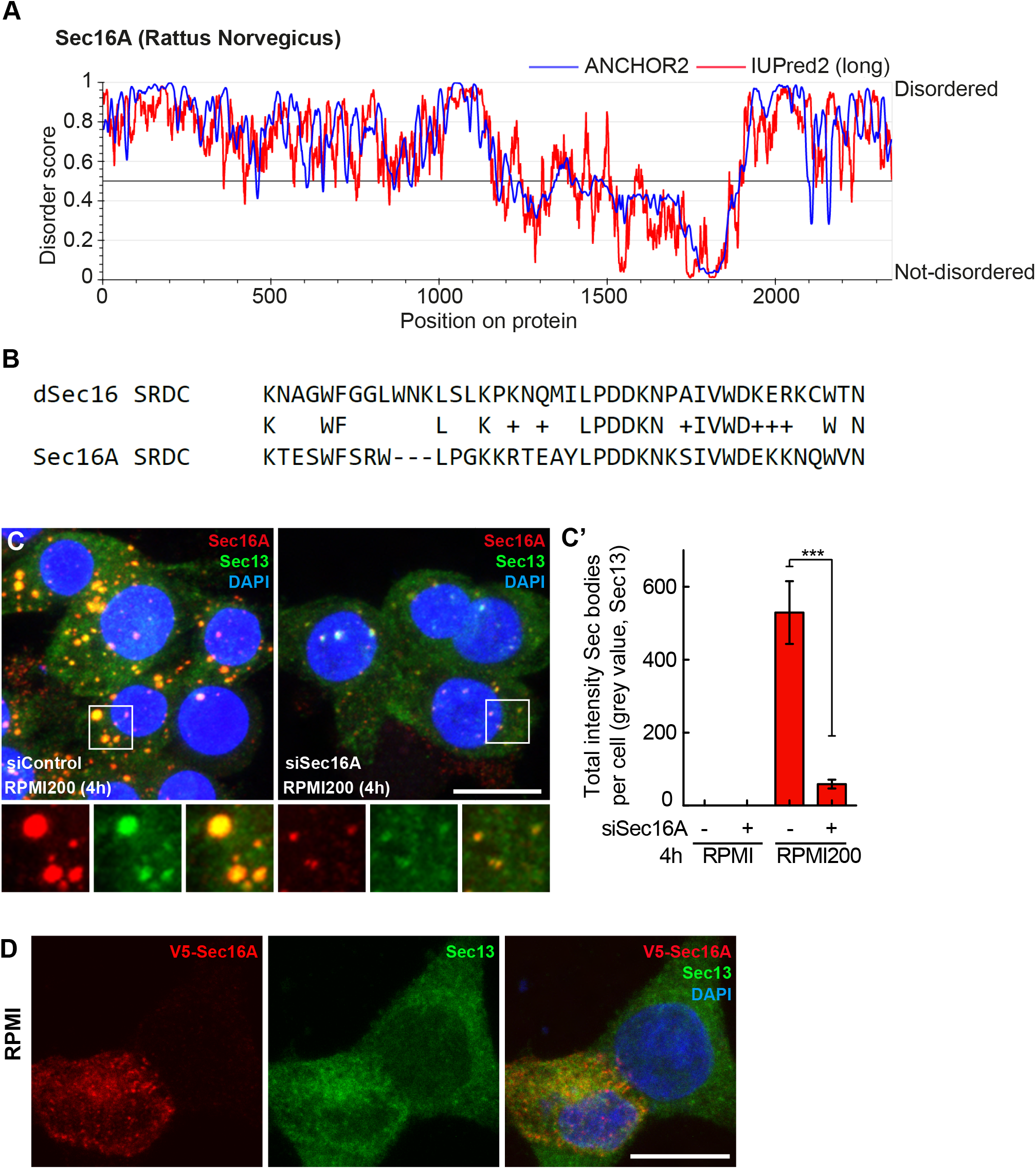
Additional information on Sec16A. **A:** Disordered regions in rat Sec16A as predicted by the database https://iupred2a.elte.hu/. A disorder score of 0.5 or higher is predicted as a disordered protein sequence. **B:** SRDC in rat Sec16A. Top panel displays the reported SRDC region in *Drosophila* Sec16 (upper row) (Aguilera-Gomez et al., 2016) which is conserved in rat Sec16A (lower row). The overlap was found by protein sequence blasting using NCBI database. The SRCD is absent in rat Sec16B. **C, C’:** Immunofluorescence visualization of Sec16A and Sec13 in Sec16A depleted INS-1 cells incubated in RPMI200 for 4 h. Note that Sec body formation (marked by Sec13) is inhibited upon Sec16A depletion. Quantification of the total intensity of Sec bodies per cell (C’); n= 24 of 2 independent experiments. **D**: Representative images of V5-Sec16A in INS-1 transfected cells cultured in RPMI medium. Cells were labeled for V5-tag, DAPI, and for endogenous Sec13 Scale bar: 10 μm (C) and (D). Error bar: SEM (C’); ***p<0.001.

**Figure 5—figure supplement 1.**
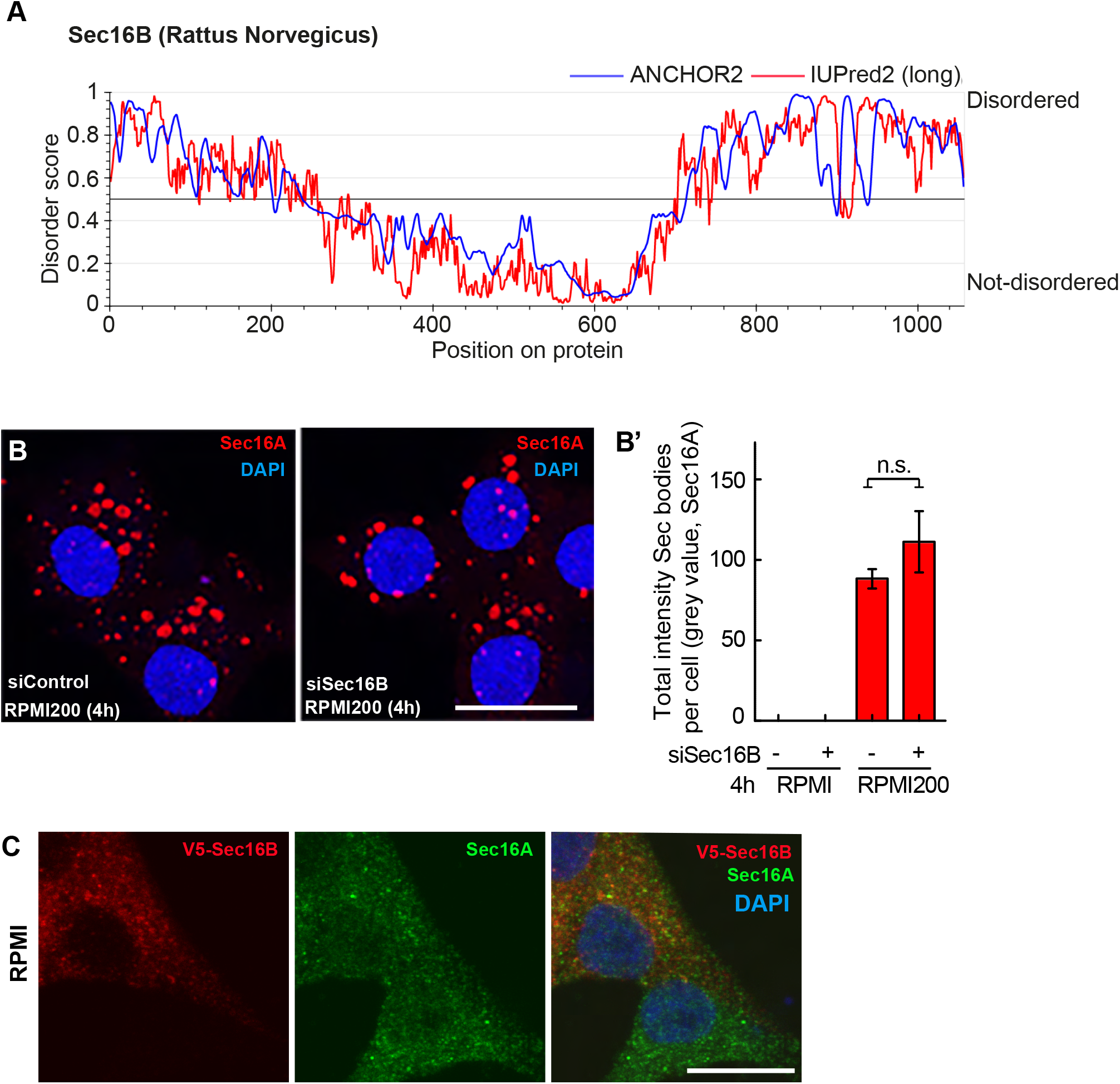
Additional information on Sec16B. **A:** Disordered regions in rat Sec16B predicted by the database https://iupred2a.elte.hu/. A disorder score of 0.5 or higher is predicted as a disordered protein sequence. **B, B’:** Immunofluorescence visualization of Sec16A in Sec16B depleted INS-1 cells incubated in RPMI200 for 4 h (B). Note that Sec body formation (marked by Sec16A) is not inhibited upon Sec16B depletion. Quantification of the total intensity of Sec bodies per cell (C’); n=25 cells from two independent experiments. **C:** Representative images of V5-Sec16B in INS-1 transfected cells cultured in RPMI medium. Cells were labeled for V5-tag, DAPI, and for endogenous Sec16A. Scale bar: 10 μm (B, C). Error bar: SEM; ns – not significant

**Figure 6—figure supplement 1.**
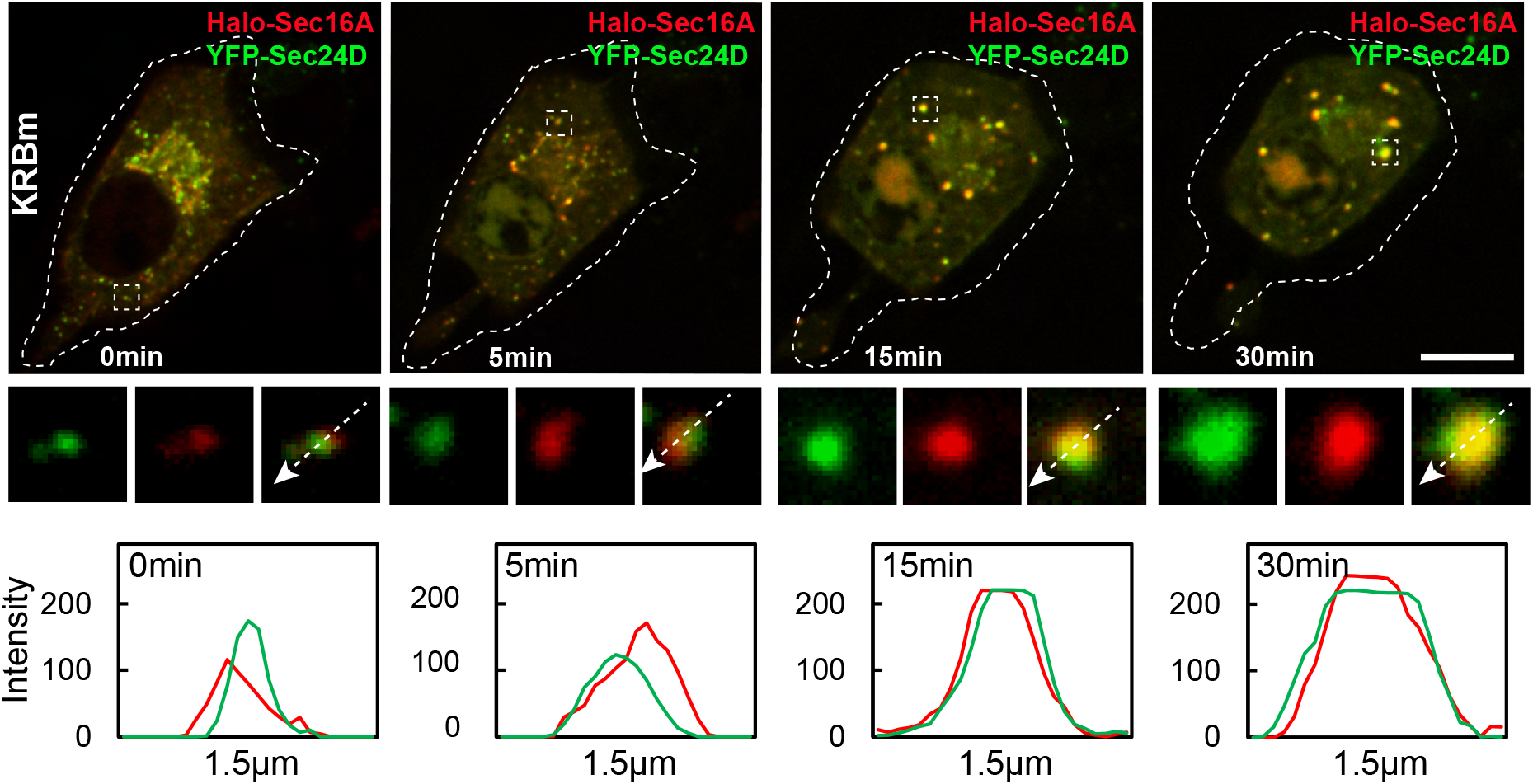
ERES components are co-recruited into newly formed Sec bodies. Representative still images of time-points 0, 5, 15 and 30 min from a live cell expressing Halo-Sec16A and YFP-Sec24D and treated with KRBm during live-cell imaging. Cells were pre-incubated with the permeable Halo-646 dye prior to imaging. Single and merged channels of selected ERES and Sec body structures are shown, and intensity profile lines displaying co-distribution of Sec16A and Sec24D. See also Video S3. Scale bar: 5 μm

**Figure 7—figure supplement 1.**
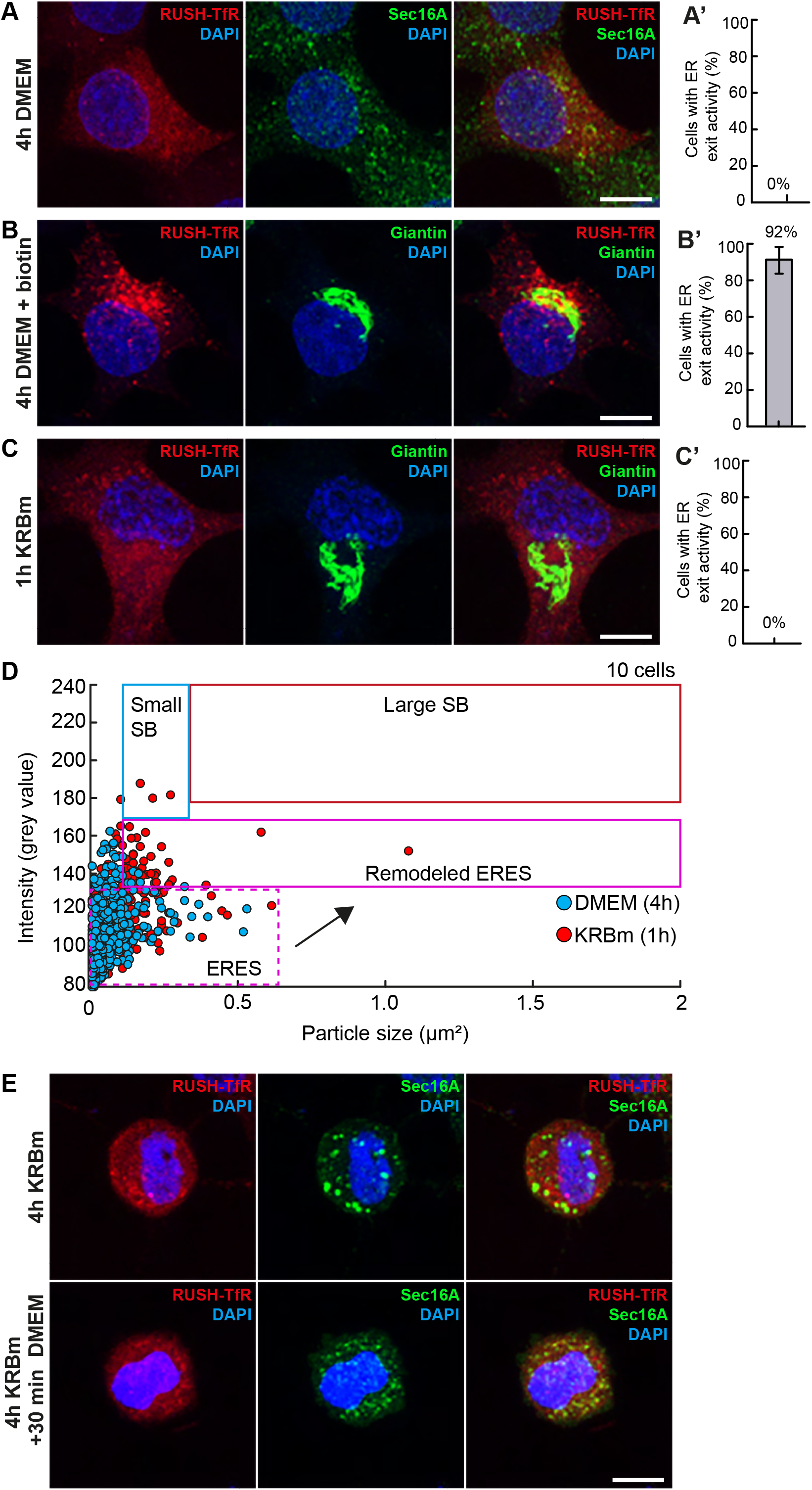
Without addition of biotin RUSH-TfR is retained at the ER A, A’: Visualization of RUSH-TfR and endogenous Sec16A in cells in DMEM (4 h) without the addition of biotin. Quantification of the ER exit activity (A’); n=13 cells **B, B’:** Visualization of RUSH-TfR and Giantin in cells cultured in DMEM (4 h) upon addition of biotin (30 min, 100 μM). Quantification of the ER exit activity (B’); n=16 cells. **C, C’:** Visualization of RUSH-TfR and Giantin in cells in KRBm (1 h) without the addition of biotin. Quantification of the ER exit activity (C’); n=15 cells. **D:** Scatterplot depicting Sec16A foci size and intensity upon incubation in DMEM (blue dots) and 1 h KRBm (red dots) of one representative INS-1 cell. Note that foci become larger and more intense after 1 h of KRBm incubation (magenta box). **E:** Visualization of RUSH-TfR and endogenous Sec16A in cells incubated in KRBm for 4 h followed by 30 min or 1 h in DMEM without biotin. Note that the Sec bodies have dissolved but that RUSH-TfR is retained in the ER. Scale bar: 10 μm Error bar: SEM (A-C)

**Supplementary Table S1.**
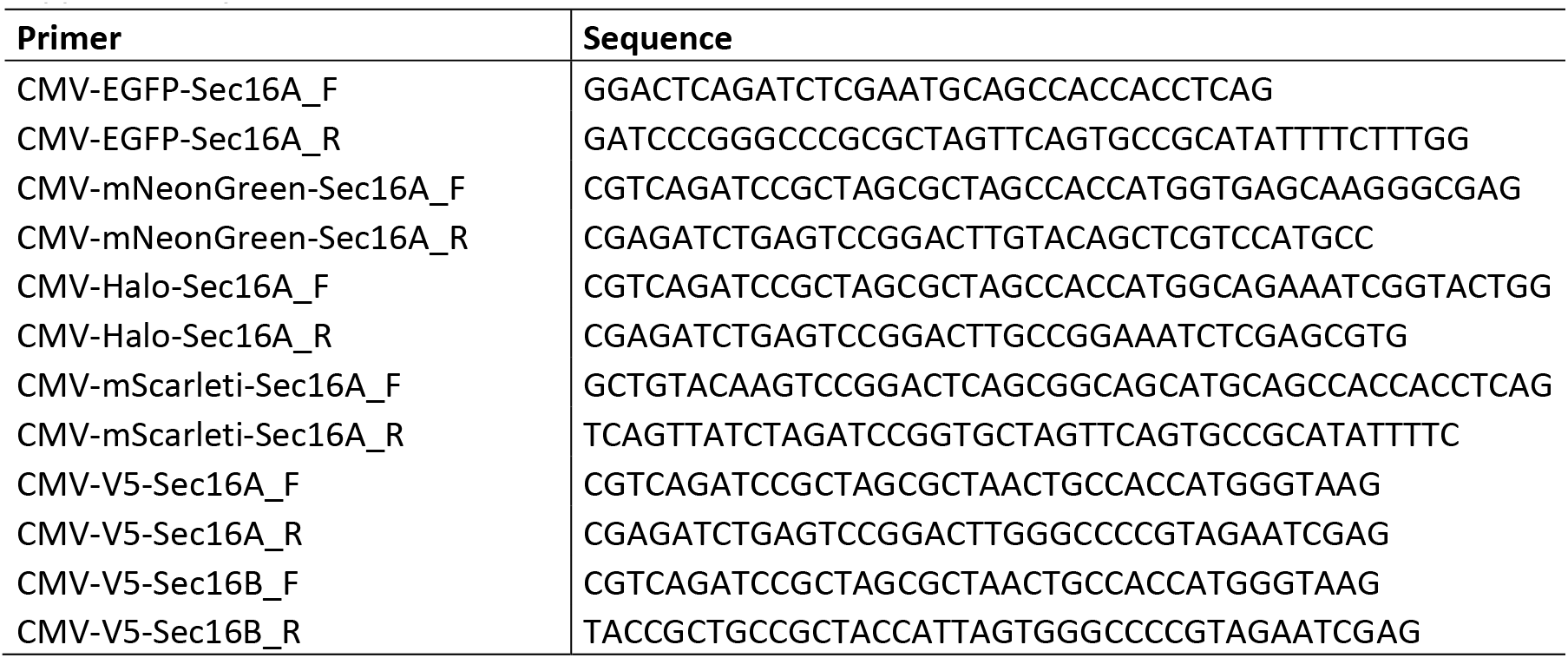

**Video S1 (related to Figure 6). Sec bodies form by fusion.** INS-1 cells expressing mNG-Sec16A were incubated with KRBm and immediately imaged every 15 seconds for 30 minutes. Right panels are zoom-in images of Sec bodies undergoing fusions at indicated time points.

**Video S2 (related to Figure 6). Simultaneously recruitment of Sec16A and Sec16B into newly formed Sec bodies**. INS-1 cells co-expressing mScarlet-Sec16A and GFP-Sec16B were incubated with KRBm and immediately imaged every 30 seconds for 30 minutes. Notice co-distribution of proteins in remodeled structures undergoing fusion events.

**Video S3 (related to Figure 6). Simultaneously recruitment of Sec16A and Sec24D into newly formed Sec bodies.** Sec bodies in INS-1 cells co-transfected with Halo-Sec16A and YFP-Sec24D. Cells were pre-incubated with a permeable Halo-646 dye prior KRB treatment during imaging every 30 seconds for 30 minutes. Notice co-distribution of proteins in remodeled structures undergoing fusion events.

## Notes

### Competing Interest Statement

The authors have declared no competing interest.

